# Evolution of genome structure in the *Drosophila simulans* species complex

**DOI:** 10.1101/2020.02.27.968743

**Authors:** Mahul Chakraborty, Ching-Ho Chang, Danielle E. Khost, Jeffrey Vedanayagam, Jeffrey R. Adrion, Yi Liao, Kristi L. Montooth, Colin D. Meiklejohn, Amanda M. Larracuente, J.J. Emerson

**Affiliations:** Department of Ecology and Evolutionary Biology, University of California Irvine, Irvine, CA 92697; Department of Biology, University of Rochester, Rochester, NY 14627; Division of Basic Sciences, Fred Hutchinson Cancer Research Center, Seattle, WA 98109; FAS Informatics and Scientific Applications, Harvard University, Cambridge, MA 02138; Department of Developmental Biology, Memorial Sloan-Kettering Cancer Center, New York, NY, 10065; Institute of Ecology and Evolution, University of Oregon, Eugene, Oregon 97403; School of Biological Sciences, University of Nebraska-Lincoln, Nebraska 68502

## Abstract

The rapid evolution of repetitive DNA sequences, including satellite DNA, tandem duplications, and transposable elements, underlies phenotypic evolution and contributes to hybrid incompatibilities between species. However, repetitive genomic regions are fragmented and misassembled in most contemporary genome assemblies. We generated highly contiguous *de novo* reference genomes for the *Drosophila simulans* species complex (*D. simulans, D. mauritiana*, and *D. sechellia*), which speciated ∼250,000 years ago. Our assemblies are comparable in contiguity and accuracy to the current *D. melanogaster* genome, allowing us to directly compare repetitive sequences between these four species. We find that at least 15% of the *D. simulans* complex species genomes fail to align uniquely to *D. melanogaster* due to structural divergence—twice the number of single-nucleotide substitutions. We also find rapid turnover of satellite DNA and extensive structural divergence in heterochromatic regions, while the euchromatic gene content is mostly conserved. Despite the overall preservation of gene synteny, euchromatin in each species has been shaped by clade and species-specific inversions, transposable elements, expansions and contractions of satellite and tRNA tandem arrays, and gene duplications. We also find rapid divergence among Y-linked genes, including copy number variation and recent gene duplications from autosomes. Our assemblies provide a valuable resource for studying genome evolution and its consequences for phenotypic evolution in these genetic model species.

## INTRODUCTION

Repetitive DNA sequences comprise a substantial fraction of the genomes of multicellular eukaryotes, occupying >40% of human and *Drosophila melanogaster* genomes (Britten and Kohne 1968; Treangen and Salzberg 2011; Hoskins et al. 2015). These sequences include repeated tandem arrays of non-coding sequences like satellite DNAs, self-replicating selfish elements like transposable elements (TEs), and duplications of otherwise unique sequences, including genes (Britten and Kohne 1968). Despite being historically considered nonfunctional, repetitive sequences are now known to play significant roles in both cellular and evolutionary processes. In many eukaryotes, satellite DNA, tandem repeats, and/or TEs constitute structures essential for genome organization and function, like centromeres and telomeres (Moyzis et al. 1988; Mason et al. 2008; Klein and O’Neill 2018; Chang et al. 2019; Hartley and O’Neill 2019). Short tandem repeats near protein-coding genes can regulate gene expression by recruiting transcription factors (Rockman and Wray 2002; Gemayel, et al. 2010), and euchromatic satellite repeats contribute to X Chromosome recognition during dosage compensation in *Drosophila* males (Menon and Meller 2012; Menon et al. 2014).

In both humans and fruit flies, genetic polymorphism composed of repetitive sequences makes up a larger proportion of the genome than all single nucleotide variants (SNVs) combined (Chakraborty et al. 2018; The 1000 Genomes Project Consortium 2015). Moreover, repetitive sequence variants can have significant fitness effects, underlie ecological adaptations, drive genome evolution, and participate in genomic conflicts (e.g. (Daborn et al. 2002; Aminetzach et al. 2005; Montchamp-Moreau et al. 2006; Tao et al. 2007b, 2007a; Fishman and Saunders 2008; Larracuente and Presgraves 2012; Ellison and Bachtrog 2013; Van’t Hof et al. 2016; Battlay et al. 2018; Chakraborty et al. 2018, 2019)). The selfish proliferation of repetitive sequences can alter protein coding genes (Lipatov et al. 2005), create intragenomic conflicts (Doolittle and Sapienza 1980; Orgel and Crick 1980) and trigger evolutionary arms races within and between genomes (Werren et al. 1988; Aravin et al. 2007; Ellis et al. 2011; Cocquet et al. 2012; Lindholm et al. 2016; Blumenstiel 2019; Parhad and Theurkauf 2019; Rathje et al. 2019). For example, centromeric repeats can drive through female meiosis, causing rapid evolution of centromere proteins to restore equal segregation (Henikoff et al. 2001). Repeats can also be the target of selfish meiotic drivers in males (e.g. (Larracuente and Presgraves 2012)), which may drive the rapid evolution of these repeats to escape the driver (e.g. (Cabot et al. 1993; Larracuente 2014)). The lack of recombination and male-limited transmission of Y chromosomes also create opportunities for conflicts involving repetitive DNA to evolve, such as sex-chromosome meiotic drive. Such conflicts have driven the proliferation of sex-linked gene families in mammals and *Drosophila* (Cocquet et al. 2012; Kruger et al. 2019)(reviewed in (Jaenike 2001)). These conflicts may also impose selection pressures that trigger the rapid turnover of Y-linked repeats (Lohe and Roberts 1990; Bachtrog 2004; Larracuente and Clark 2013; Mahajan et al. 2018; Wei et al. 2018).

The very nature of repetitive sequences makes them difficult to study. Whole-genome shotgun sequencing of reads shorter than common repeats yields erroneous, fragmented, and incomplete genome assemblies in repetitive regions (Salzberg and Yorke 2005; Hoskins et al. 2002; Alkan et al. 2011; Treangen and Salzberg 2011; Hoskins et al. 2015). Reference-quality genomes have historically been available only for distantly related species, making it difficult to investigate the evolutionary dynamics of repetitive sequences (reviewed in (Plohl et al. 2012; Lower et al. 2018; Blumenstiel 2019)). Long-read based assemblies help solve these challenges because they can be nearly complete, contiguous, and accurate even in repetitive genomic regions (Steinberg et al. 2014; Berlin et al. 2015; Chaisson et al. 2015; Chakraborty et al. 2016; Mahajan et al. 2018; Chakraborty et al. 2018; Solares et al. 2018; Chang and Larracuente 2019).

To understand the contributions of repetitive sequences to genome structure and evolution, we sequenced and assembled reference-quality genomes of *Drosophila simulans, D. sechellia*, and *D. mauritiana*. These three species, collectively known as the *Drosophila simulans* species complex (or sim-complex (Kliman et al. 2000)), comprise the nearest sister species to *D. melanogaster*, and are virtually equally related to each other (Fig. 1A), likely as a consequence of rapid speciation (Garrigan et al. 2012; Pease and Hahn 2013). The four fruit fly species together comprise the *D. melanogaster* species complex (or mel-complex) (Hey and Kliman 1993). The mel-complex serves as a model system for studying speciation (Tao et al. 2001; Wu 2001; Meiklejohn et al. 2018), behavior (Ding et al. 2019), population genetics (Kliman et al. 2000; Begun et al. 2007; Garrigan et al. 2012), and molecular evolution (Moriyama and Powell 1997; Ranz et al. 2007; Hu et al. 2013). All four species are reproductively isolated from one another, producing either sterile or lethal hybrids (Barbash 2010). They exhibit unique ecological adaptations: *D. sechellia* larvae specialize on a host fruit toxic to most other *Drosophila* species (R’Kha et al. 1991) whereas *D. melanogaster* larvae can thrive in ethanol concentrations lethal to the sim-complex species (Merçot et al. 1994). In euchromatic regions, these species show ∼95% sequence identity (Begun et al. 2007; Garrigan et al. 2012). However, the degree of interspecific divergence in repetitive genomic regions that are not represented in current assemblies is unknown (Chakraborty et al. 2018; Miller et al. 2018).

**Figure 1.**
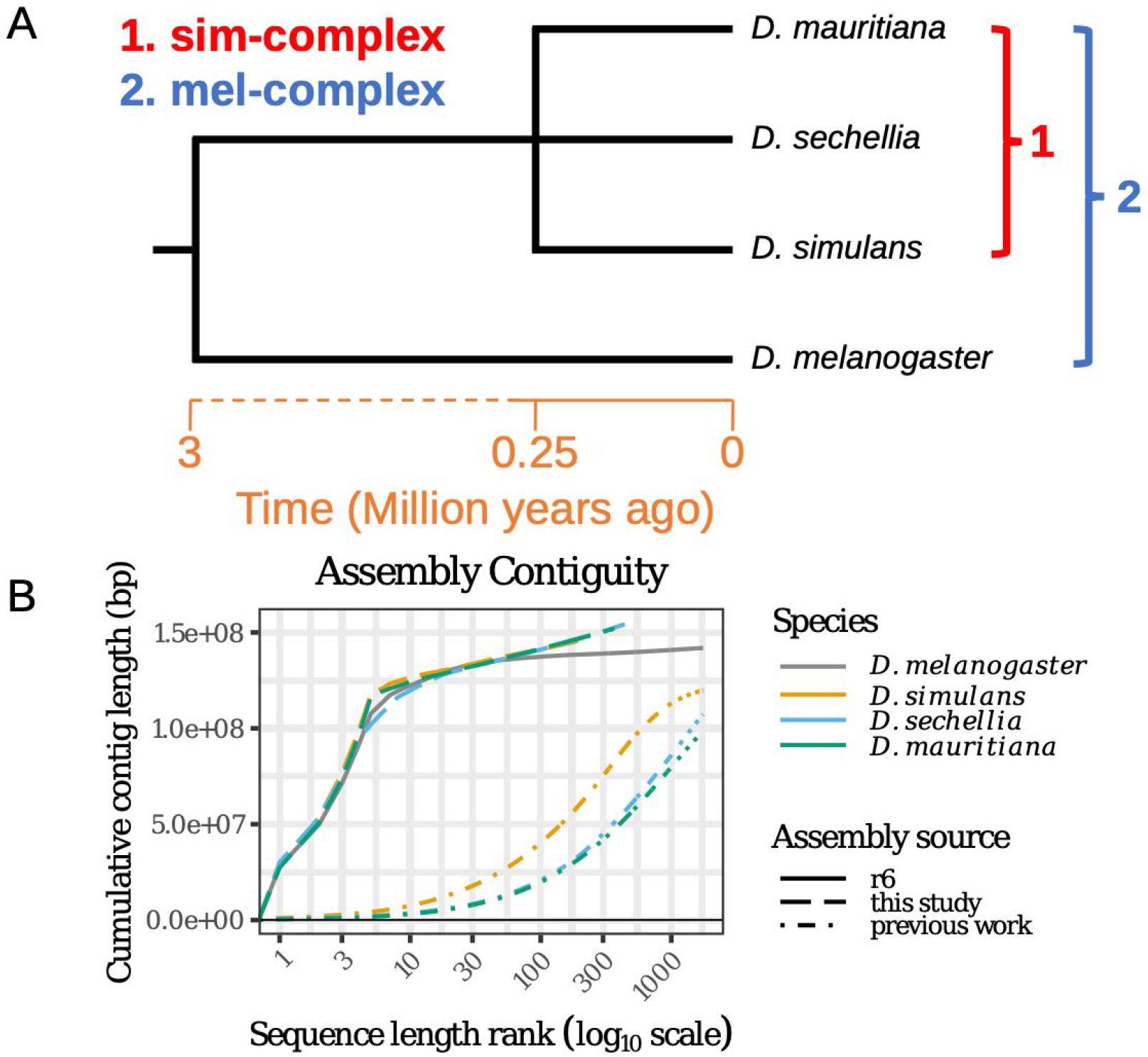
Reference-quality *de novo* genome assemblies of the *Drosophila melanogaster* species complex. (A) Phylogeny showing the evolutionary relationship among the members of four mel-complex species. (B) Contiguities of the new assemblies from the sim-complex and the reference assembly of *D. melanogaster* (R6). The contigs were ranked by their lengths and their cumulative lengths were plotted on the Y-axis. The colors represent different species. The *D. melanogaster* genome is the release 6 assembly (Hoskins et al. 2015). For previous work, *D. simulans* is ASM75419v3 (Hu et al. 2013), *D. sechellia* (r1.3) is from (*Drosophila* 12 Genomes Consortium 2007), and *D. mauritiana* is from (Garrigan et al. 2014).

Here we use high-coverage long read sequencing to assemble sim-complex genomes *de novo*, permitting us to resolve repetitive regions that have, until now, evaded scrutiny. These assemblies are comparable in completeness and contiguity to the latest release of the *D. melanogaster* reference genome. Our results uncover a dynamic picture of repetitive sequence evolution that leads to extensive genome variation over short timescales.

## RESULTS

### Contiguous, accurate, and complete assemblies resolve previous misassemblies

We collected deep (100–150 fold autosomal coverage) long read sequence data from adult male flies (supplemental Fig. S1–2; supplemental Table S1) to assemble reference-quality genomes *de novo* for the three sim-complex species. Our assemblies are as contiguous as the *D. melanogaster* reference (Hoskins et al. 2015) (Fig. 1B, supplemental Fig. S3, supplemental Table S2). In all three species, single contigs span the majority of each chromosome arm, except the X Chromosome in *D. sechellia*. Our scaffolds include the entirety of the euchromatin and large stretches of pericentric heterochromatin (Fig. 1B, Fig. 2, supplemental Fig. S4). We assembled more than 20-Mbp of pericentric heterochromatin (Fig. 2A), overcoming difficulties associated with these genomic regions (Khost et al. 2017; Chang et al. 2019).

**Figure 2.**
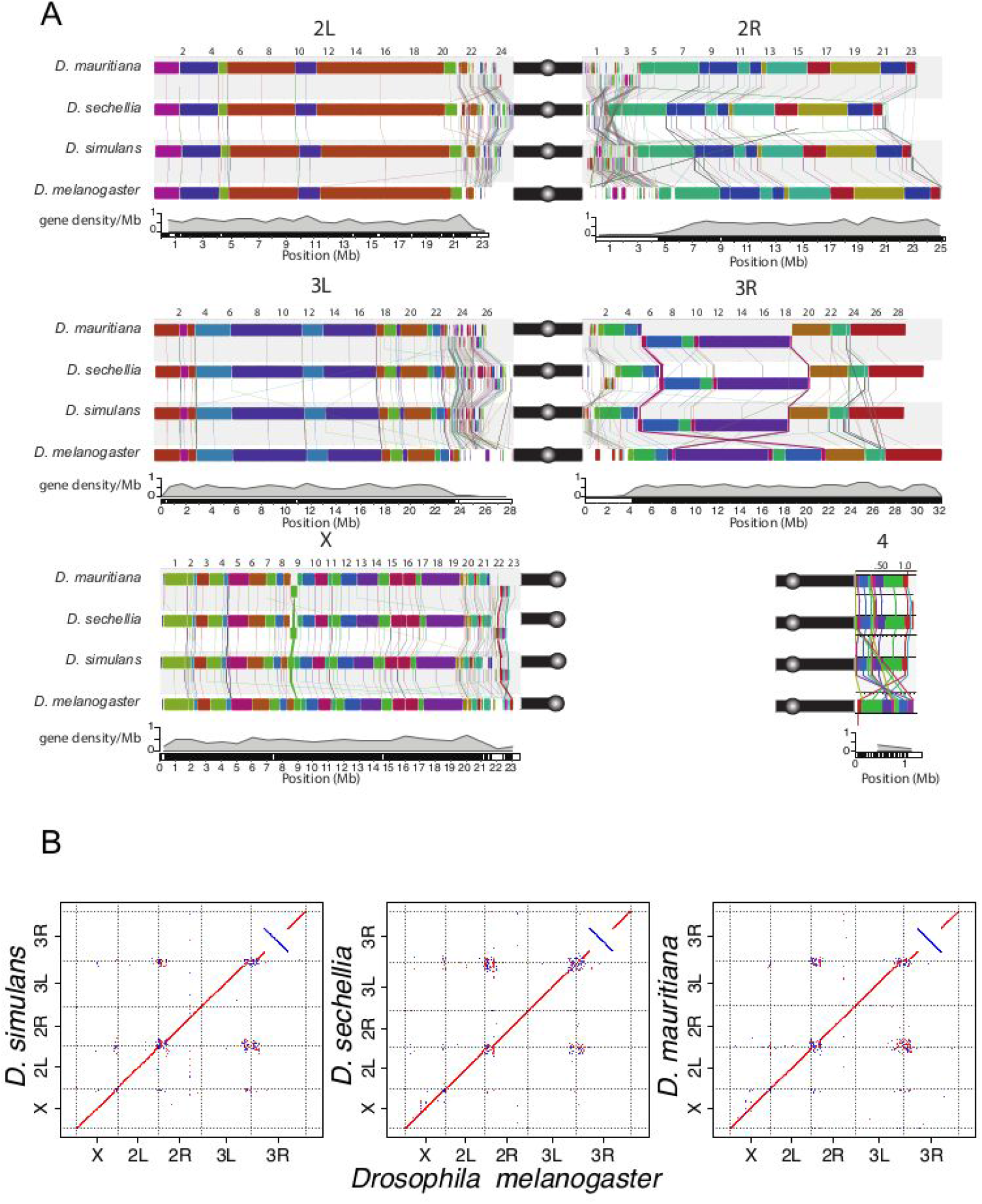
Chromosomal rearrangements in the sim-complex species. We used mauve (A) and minimap2 (Li 2018) (B) to show synteny between the members of the sim-complex and *D. melanogaster*. A) Colored rectangles show positions of syntenic collinear blocks free from internal rearrangements compared to the *D. melanogaster* reference (r6; see details in Materials and Methods). Each chromosome arm is plotted with its own scale, with position in megabases indicated above each chromosome. Blocks that appear below the black line are in an inverse orientation. Lines connect homologous colored blocks between genomes, and crossing lines indicate structural rearrangements. Along the euchromatic chromosome arms, there are three major inversion events (X, 3R, and 4). The heterochromatic regions have significantly more rearrangements than the euchromatin (see text). Pericentromeric heterochromatic regions are marked with a solid black bar and the circles correspond to centromeres. B) The dot plots for the whole genome and each chromosome arm between the sim-complex species and *D. melanogaster*.

Comparison of our assemblies to the *D. melanogaster* genome recovers synteny expected between the species across major chromosome arms (Fig. 2; supplemental Fig. S4). Genome-wide, ∼15% of sim-complex genome content fails to align uniquely to *D. melanogaster*. Within aligned sequence blocks, the sim-complex species show ∼7% divergence from *D. melanogaster* (supplemental Fig. S5). Preservation of synteny between the genomes suggests that there are no large errors, which is further supported by the evenly distributed long-read coverage (supplemental Fig. S1–2) and mapping of BAC sequences across the assembled chromosomes (supplemental Fig. S6; supplementary text). We corrected errors previously noted in the draft assemblies of these species (supplemental Fig. S7 and supplemental Table S3), including a ∼350-kb 3L subtelomeric fragment misassembled onto the 2R scaffold in the previous *D. simulans* assembly (Schaeffer et al. 2008). Our assemblies are also highly accurate at the nucleotide level, as concordance between our assemblies and Illumina data is comparable to that of *D. melanogaster* (cf QV = 44.0–46.3 for sim-complex species vs. 44.3 for *D. melanogaster*; supplemental Table S4). The sim-complex assemblies are highly complete, with numbers of single-copy conserved Dipteran orthologs (BUSCO; (Simão et al. 2015)) comparable to that of *D. melanogaster* (98.6-99% BUSCO; supplemental Table S5). Moreover, we detected more *D. melanogaster* orthologous genes in our sim-complex assemblies compared to the previous assemblies (supplemental Table S6; supplementary text).

We also assembled entire *Wolbachia* genomes from *D. mauritiana* (*w*Mau) and *D. sechellia* (*w*Sech) (supplemental Table S7); our *D. simulans w*^XD1^ strain was not infected with *Wolbachia*. Our assemblies reveal extensive and previously unknown structural divergence between closely related *Wolbachia* genomes. *w*Sech is 95.1% identical to *w*Ha (supergroup A) from *D. simulans*. We detect a single inversion differentiating *w*Sech from *w*Ha (supplemental Fig. S8A). *w*Mau is 95.8% identical to *w*No from *D. simulans* (supergroup B), and >99.9% identical to other recently published *Wolbachia* genomes from *D. mauritiana* (available from NCBI under accessions CP034334 and CP034335)(Lefoulon et al. 2019). We infer extensive (15) structural rearrangement events between recently diverged *Wolbachia* lineages, *w*No and *w*Mau, under double-cut-and-join (DCJ) model (supplemental Fig. S8B)(Lin and Moret 2008). A recent study of *Wolbachia* from different isolates of *D. mauritiana* identified four deletions in *w*Mau relative to *w*No (Meany et al. 2019). Our assemblies indicate that these deletions are associated with other SVs. Three of the four deletions (CNVs 1, 3, and 4 in supplemental Fig. S8C) occur at rearrangement breakpoints, while the fourth (CNV 2) shows a segment repeated in *w*No flanking the segment deleted in *w*Mau. Finally, *w*Mau maintains a single-copy segment in one of the deletions (CNV 1) which itself is a dispersed duplication in *w*No (supplemental Fig. S8C). It remains unclear whether any of these structural changes contribute to the lack of fecundity effects or cytoplasmic incompatibility caused by infection with *w*Mau (Meany et al. 2019).

### Clade- and species-specific genomic rearrangements

We computed locally collinear alignment blocks with Mauve (Lin and Moret 2008) to infer genomic rearrangements between species. We discovered 535–542 rearrangements between *D. melanogaster* and the sim-complex (∼90 mutations per million years), and 113–177 rearrangements within the sim-complex (226–354 mutations per million years; supplemental Table S8). Heterochromatic regions harbor 95% of all genomic rearrangements (supplemental Fig. S9 and supplemental Table S8). In euchromatin, there is an enrichment of rearrangements on the X Chromosome: 63% of all identified rearrangements (17/27) between *D. melanogaster* and the sim-complex species and all but one (12/13) rearrangement within the sim-complex species are X-linked (Fig 2A; supplemental Table S8).

Within euchromatin, *D. simulans, D. mauritiana*, and *D. sechellia* differ from *D. melanogaster* by 23, 25, and 21 inversions, respectively. We recovered the 13.6-Mb *D. melanogaster*-specific 3R inversion (In(3R)84F1; 93F6–7; 3R:8,049,180–21,735,108) that was initially characterized cytologically (Sturtevant and Plunkett 1926) and confirmed by breakpoint cloning (Ranz et al. 2007)(Fig. 2). Among nine inversions shared in all sim-complex species, four are also present in the outgroup species *D. yakuba* and *D. ananassae*, suggesting that they occurred in the *D. melanogaster* lineage. The remaining five are found only in the sim-complex species. The sim-mau, sim-sec, and mau-sec species pairs share 5, 3, and 4 euchromatic inversions absent in the third species, respectively. For example, *D. sechellia* and *D. mauritiana*, but not *D. simulans*, share a 460-kb X-linked inversion (X:8,744,323–9,203,725 and X:8,736,133–9,203,526, respectively) spanning 45 protein-coding genes (Fig. 2, supplemental Fig. S10A).

We also observe evidence for two large (>100-kb) inversions within pericentromeric heterochromatin on Chromosomes 3 and X (Fig. 2, supplemental Fig. S11A–D). Because *D. erecta* shares the same configuration as the sim-complex species, the pericentric inversion on Chromosome 3 likely occurred in the *D. melanogaster* lineage (supplemental Fig. S12). We also observed an ∼700-kb inversion in the X heterochromatin of sim-complex species spanning 35 genes (22.4–23.1Mb on *D. melanogaster* X; Fig. 2, supplemental Fig. S4B, 4H, 4N and S10). This inversion is sim-complex specific, and is absent in *D. melanogaster, D. yakuba*, and *D. erecta*. We also find large, species-specific heterochromatic inversions on 3R in *D. sechellia* (Fig. 2, supplemental Fig. S11A–B) and 2R in *D. mauritiana* (supplemental Fig. S13).

### Repetitive DNA

Our annotations of repetitive DNA (supplementary File S1) revealed substantially greater repeat abundance in the sim-complex genomes compared to older assemblies of these species (supplemental Fig. S14). On the five large chromosome arms, the density of repetitive elements increases approaching the euchromatin-heterochromatin boundary, consistent with patterns of TE density in *D. melanogaster* (Kaminker et al. 2002; Bergman et al. 2006)(Fig. 3). Below we describe our analyses of the different classes of repetitive elements.

**Figure 3.**
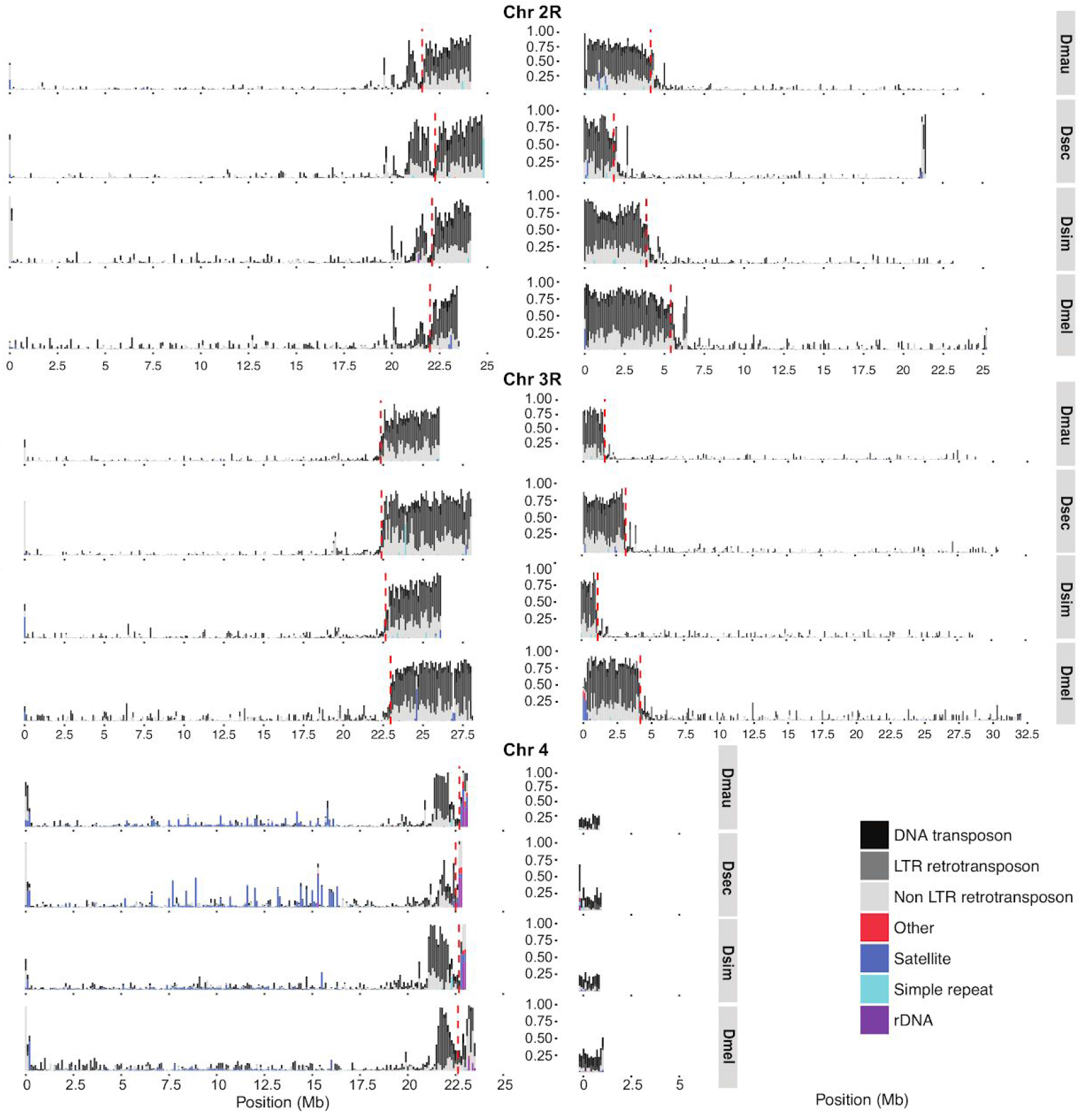
The repeat content across the chromosome arms in mel-complex species. We estimated the repeat content in the genome using RepeatMasker (Smit et al. 2013). Each bar represents the proportion of different repeat types in 100-kb windows. The red dashed vertical lines indicate the euchromatin-heterochromatin boundaries.

#### Distribution of satellites

We identified three novel complex satellite arrays in the sim-complex, which we named for their monomer size (90U, 193XP, and 500U). 500U is located primarily on the unassigned contigs and cytologically near centromeres (Chang et al. 2019; Talbert et al. 2018). The 90U satellite corresponds to one of the non-transcribed ribosomal DNA (rDNA) spacer (NTS) subunits (Stage and Eickbush 2007). 90U repeats are adjacent to the 28S rDNA subunit and the 240-bp NTS repeat sequences, both on X-linked and unassigned contigs. We find a large 193XP locus in the pericentromeric heterochromatin adjacent to, but distinct from, the rDNA locus. In *D. simulans* and *D. mauritiana*, the 193XP loci span at least 48 kb. The 193XP locus is shared across the sim-complex but is absent in the outgroup species *D. melanogaster, D. erecta*, and *D. yakuba*, suggesting that it arose in the ancestor of the sim-complex. Consistent with our assemblies, we detect fluorescence *in situ* hybridization signal for 193XP only on the X pericentromere in the sim-complex (supplemental Fig. S15).

We also find smaller satellite arrays in the euchromatin (supplemental Table S9) as has been previously reported (Kuhn et al. 2012; Gallach 2014; Waring and Pollack 1987; DiBartolomeis et al. 1992). Satellites comprise only ∼0.07% of bases in autosomal euchromatin, but they comprise 1% of X-linked euchromatin in *D. melanogaster* and *D. simulans*, up to 2.4% in *D. mauritiana*, and more than 3.4% in *D. sechellia* (supplemental Table S9). The number in *D. sechellia* is a minimum estimate because its assembly contains 6 gaps in X-linked euchromatic satellite regions. The location, abundance, and composition of euchromatic satellites differ substantially between species. For example, a complex satellite called *Rsp-like* (Larracuente 2014) recently expanded in *D. simulans* and *D. mauritiana*, and inserted into new X-linked euchromatic locations within existing arrays of another satellite called *1*.*688*. Large blocks of *1*.*688 (Lohe and Brutlag 1987)* and *Rsp-like* (Larracuente 2014; Sproul et al. 2020) differ in abundance and location in the heterochromatin of all four species.

#### Transposable elements

We annotated euchromatic TEs across *D. melanogaster* and the three sim-complex species (see Methods). Unless otherwise noted, our results are based on comparisons of TE content (i.e. number of bases) rather than the number of TE insertions (i.e. number of events). We find that the sim-complex genomes host 67–83% as much TE sequence as *D. melanogaster* (Fig. 4A). The major difference in TE composition among the four mel-complex species is the enrichment of LTR retrotransposons in *D. melanogaster* (Kaminker et al. 2002; Bergman and Bensasson 2007; Kofler et al. 2015), which carries 1.3–1.8 Mbp more LTR bases than the three sim-complex species (Fig. 4A–B). Both DNA and non-LTR transposon content in *D. melanogaster* are similar to those of the sim-complex species (Fig. 4A-B). Most TE bases (66–72%) in the sim-complex are found in only one species’ genome (Fig 4C), implying that these sequences have resulted from recent transposon activity.

**Figure 4.**
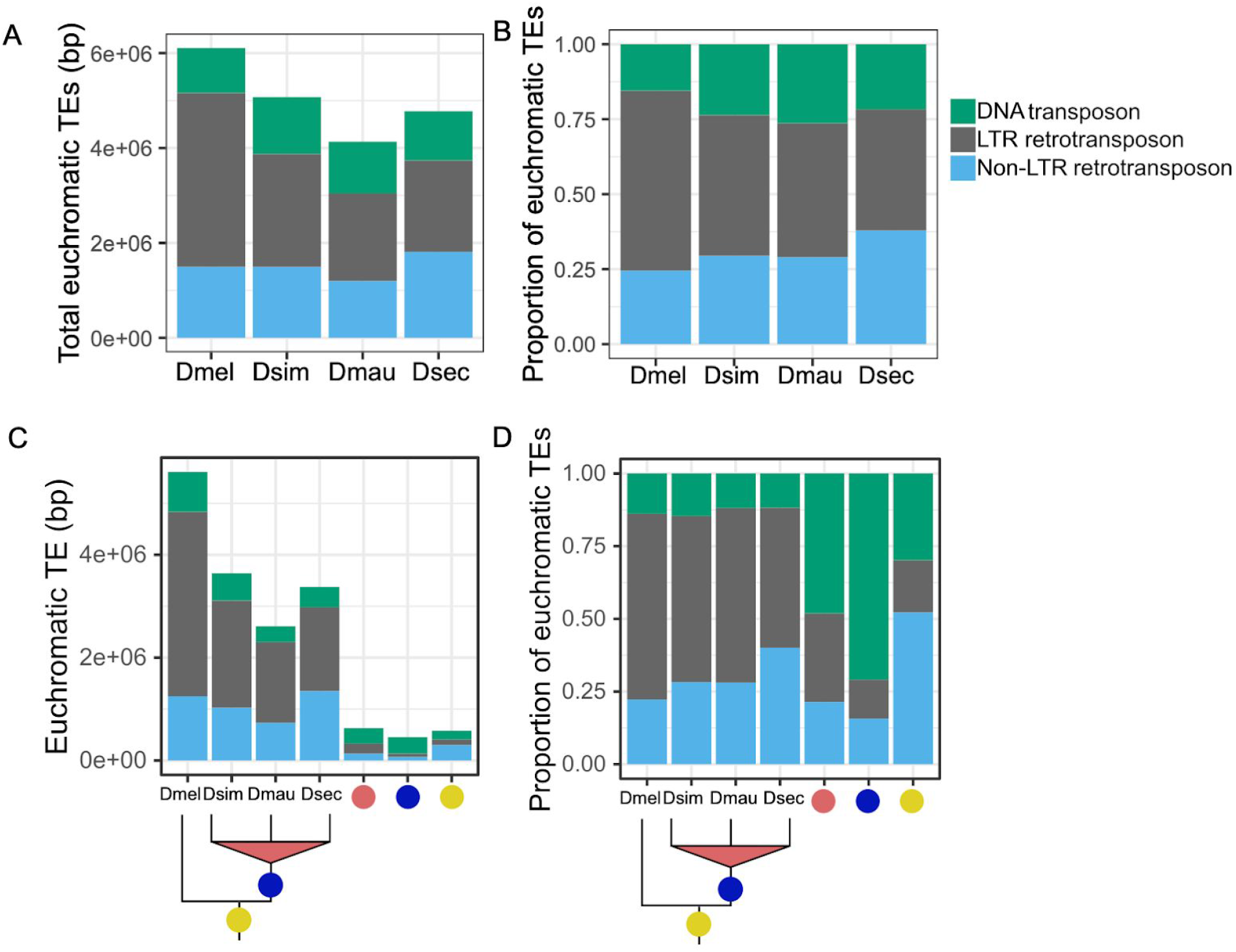
Euchromatic transposon sequence content in each species and their ancestral lineages in the mel-complex. The bars represent the absolute content (A & C) or relative proportion within each category (B & D) of TE bases due to DNA, and LTR and non-LTR retrotransposon TEs. A and B show total TE content in each species. Panels C and D show the TE content confined to specific lineages. In panels C and D the species names indicate TE sequence found only in that genome; the red circles indicate TE content found in two sim-complex species; the blue circles indicate TEs found in the sim-complex but not *D. melanogaster*; and the yellow circles indicate TE sequence found in all four mel-complex species.

We also find that TE composition differs across the lineages that gave rise to these four species (Fig. 4D; supplemental Fig. S16). Within the syntenic TE content shared by all four mel-complex species, non-LTR retrotransposon sequence is the most prevalent (52%), followed by DNA transposons (30%) and LTR retrotransposons (18%). In contrast, orthologous TE sequences present in all three sim-complex species but not *D. melanogaster* are enriched in DNA transposons, which make up 71% of this orthologous sequence (Fig. 4D) despite being shorter than other TE classes (supplemental Fig. S17). The INE-1 element (also called DINE-1 or DNAREP1) is a highly abundant DNA transposon in *Drosophila* (Quesneville et al. 2005; Yang and Barbash 2008) that has contributed to an abundance of shared INE-1 elements fixed in mel-complex (Sackton et al. 2009). In our assemblies, INE-1 makes up 46% of shared TE content in the lineage leading to the sim-complex, and a significant, but smaller proportion (13.7%) in the mel-complex lineage. The TE composition of species-specific sequences is dominated by LTR elements (48–57%) followed by non-LTR elements (27–40%), with a smaller contribution of DNA elements (12–16%) (Fig. 4D).

TE sequences can get incorporated into host genes (Lipatov et al. 2005). We find 0.8–1.6 Mb of TE sequence that overlaps with gene models in *D. melanogaster* and *D. simulans*. A small minority of young genic TEs (present only in *D. melanogaster*, only in *D. simulans*, or in the sim-complex but not *D. melanogaster*) are exonic (7–18%; Table 1). In contrast, half of the TE sequence present in all four mel-complex species is exonic (52%). This preponderance of exonic TE content in the mel-complex ancestor exceeds even the enrichment of non-LTR sequence across the whole genome (Compare supplemental Fig. S18 and Fig. 4D).

**Table 1.**
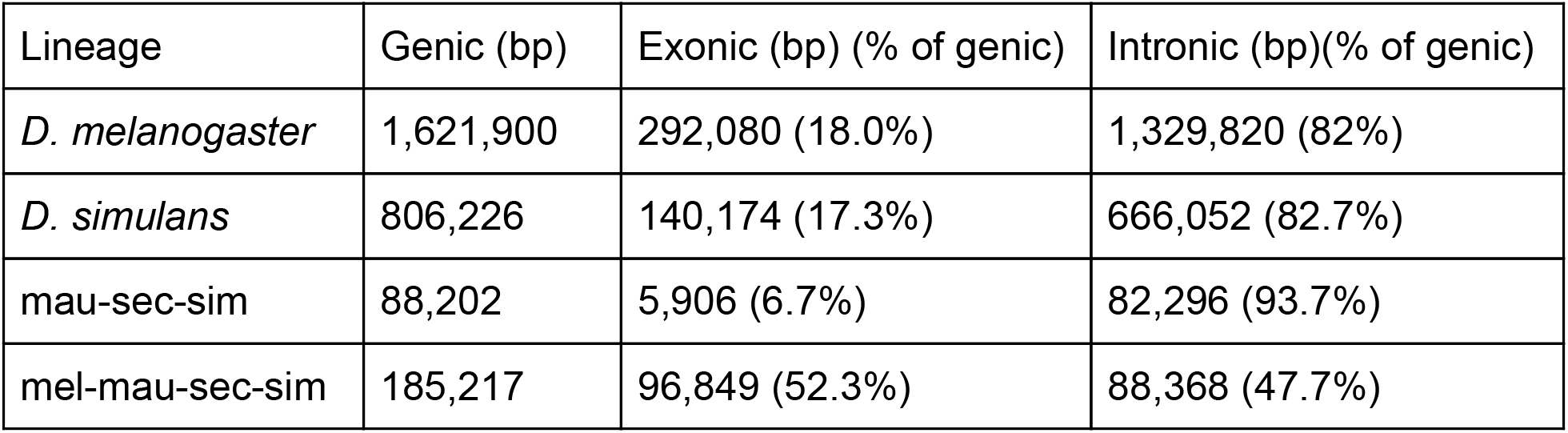
TE bases in *D. melanogaster, D. simulans*, the ancestral lineages of the sim-complex species (mau-sec-sim) and the mel-complex species (mel-mau-sec-sim) in 6,984 conserved genes. Verification of transcript expression in the sim-complex is based on Iso-Seq from *D. simulans* (Nouhaud 2018), so species-specific classifications are not available for *D. mauritiana* or *D. sechellia*.

### Intron indel mutation patterns

We compared 21,860 introns in 6,289 orthologous genes with conserved annotation positions in all four mel-complex species. We find that introns containing TE-derived sequences or complex satellites (‘complex introns’) range from 530–850bp longer in *D. melanogaster* (paired t-tests, all p-values < 0.001; supplemental Fig. S19), due largely to longer intronic TEs (mean TE length = 4,132 bp) compared to the sim-complex species (mean TE lengths of *D. simulans* = 2,429 bp, *D. mauritiana* = 2,253 bp, *D. sechellia* = 2,287 bp; supplemental Fig. S20). Among sim-complex species, *D. sechellia* has the longest complex introns in heterochromatin (both paired t-tests p-values < 0.05) but not in euchromatin (paired t-tests p-value > 0.09; supplemental Table S10 and supplemental Fig. S19). Similar to the complex introns, introns without transposons or complex satellite sequences (‘simple introns’) are significantly longer in *D. melanogaster* than the sim-complex species (paired t-tests, all p-values < 1e-7, supplemental Fig. S19; supplemental Table S10), but the mean length difference is less than 3 bp (supplemental Table S10). Consistent with a previous report (Presgraves 2006), we infer that this difference is partly due to an insertion bias in *D. melanogaster* (see supplementary text).

### Tandem duplication

We found 97 euchromatic tandem duplications shared by all three sim-complex species but absent from *D. melanogaster* (supplemental Table S11). Among these, at most 11 overlapped with duplications observed in the outgroup *D. yakuba*, suggesting that at least 86 duplications originated during the ∼2.5 million years in the ancestral lineage of the sim-complex since diverging from *D. melanogaster*. Of these duplications, 72% (62/86) overlap exons, 37% (32/86) overlap complete protein-coding sequence, and 15% (13/86) overlap one or more full-length *D. melanogaster* genes. In total, 32 complete coding sequences were duplicated, or 12.8 new genes / million years. Similar to the polymorphic duplicates in *D. simulans* (Rogers et al. 2014), tandem duplications fixed in the sim-complex ancestral lineage are strongly enriched on the X Chromosome relative to the autosomes (43/86; p-value < 1×10^−10^, proportion test against X-linked genes as a proportion of all genes, or 0.158). As a result, the X Chromosome carries both an excess of duplicates spanning full coding sequences (15 X-linked, 17 autosomal; p-value = 4.7×10^−6^, proportion test against the proportion of X-linked genes) as well as full transcripts (6 X-linked, 7 autosomal; p-value = 2.8×10^−3^, proportion test against the the proportion of X-linked genes).

Several duplication events include genes associated with divergence of important phenotypes, including: spermatogenesis (*nsr*; *(Ding et al. 2010)*), meiosis (*cona*), odorant binding (*obp18a*), chromosome organization (*HP1D3csd*), and behavior (*RhoGAP18B*)(Rothenfluh et al. 2006). Many are absent in the previous assemblies of the sim-complex species. For example, we discovered a new X-linked 3,324-bp duplication that copied the genes *maternal haploid* (*mh*) and *Alg14*. Analysis of *D. mauritiana* and *D. simulans* RNA-seq reads from our strains and Iso-Seq reads from another *D. simulans* strain (Nouhaud 2018) suggests that the distal copy (*mh-d*) produces a shortened transcript and protein compared to *mh-p* and the ancestral *mh* (Fig. 5A and supplemental Fig. S21–22). *mh-p* has female-biased expression in *D. simulans*, as does *mh* in *D. melanogaster*, where it has an essential maternal effect in zygotic cell division (Loppin et al. 2001; Delabaere et al. 2014). In contrast, *mh-d* exhibits testis-biased expression (Fig. 5B and supplemental Fig. S21), suggesting that *mh-d* may have acquired a male-specific function in the sim-complex species.

**Figure 5.**
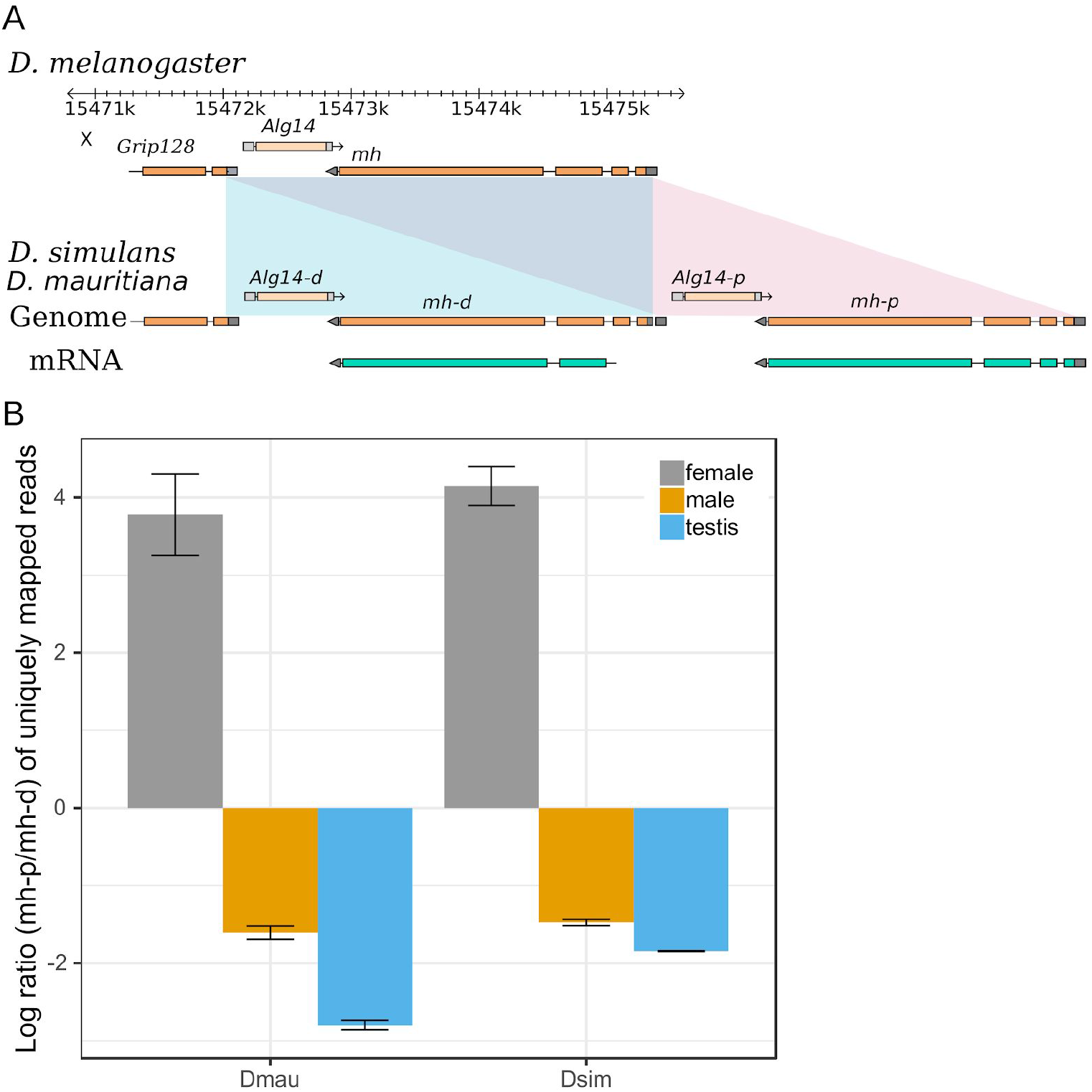
The expression divergence of *maternal haploid* (*mh*) duplicates in the sim-complex species. A) The sim-complex shares a tandem duplication of *mh* and *alg14* genes. The expression of both *mh* copies is supported by Iso-Seq and Illumina transcriptome data. B) The proximal copy of *mh* (*mh-p*) is primarily expressed in females and the distal copy (*mh-d*) shows testis-biased expression in both *D. mauritiana* and *D. simulans*.

We also uncovered a 4,654-bp tandem duplication located entirely in an inverted segment of the pericentric heterochromatin on the sim-complex X Chromosome that partially copied the gene *suppressor of forked* (*su*(*f*)) (supplemental Fig. S23). This duplicate is absent in the previous *D. mauritiana* assembly (Garrigan et al. 2014) and the reference genomes of *D. simulans* (r2.02) and *D. sechellia* (r1.3). The proximal *su(f)* copy is missing the first 12 codons but retains the rest of the ORF of the parental *su(f)* coding sequence, including the stop codon (supplemental Fig. S23).

### Evolution of tRNA clusters

Nuclear tRNAs are distributed both individually and in clusters containing identical copies coding for the same amino acids (isoacceptor tRNAs) and interspersed with those coding for different amino acids (alloacceptor tRNAs). Previous analyses found a smaller complement of tRNAs in *D. simulans* than in *D. melanogaster* (*Drosophila* 12 Genomes Consortium 2007), though it could have been due to a difference in assembly quality (*Drosophila* 12 Genomes Consortium 2007; Rogers et al. 2010; Velandia-Huerto et al. 2016). We found genome-wide tRNA counts to be similar between the species, ranging from 295 in *D. melanogaster* to 303 copies in *D. sechellia* (supplemental Fig. S24 and supplemental Table S12).

Our count of tRNAs in *D. simulans* (300 tRNAs) is substantially higher than previously reported using an older assembly (268 and 255 tRNAs; (Rogers et al. 2010; Velandia-Huerto et al. 2016), respectively), suggesting that the high rates of tRNA loss reported previously were due to assembly errors.

We identified putative tRNA orthologs using alignments encompassing tRNAs, and identified syntenic blocks of tRNAs that differed in copy-number, identity (isotype), anticodon, and pseudogene designations (Fig. 6). To confirm gains or losses, we employed a BLAST-based approach, similar to methods used by (Rogers et al. 2010), to identify regions flanking orthologous tRNA clusters. We identified four tRNA anticodon shifts, including one isoacceptor and three alloacceptor shifts (Fig. 6B), consistent with previous reports (Rogers et al. 2010; Rogers and Griffiths-Jones 2014; Velandia-Huerto et al. 2016). We did not detect a previously identified alloacceptor shift (Met CAT > Thr CGT) (Velandia-Huerto et al. 2016; Rogers and Griffiths-Jones 2014), which could be due to allelic variation within *D. simulans*. In each case, the derived tRNA sequence was otherwise similar to and retained the predicted structure of the ancestral tRNA, suggesting that the alloacceptor shifts cause the aminoacyl tRNA synthetase (aaRS) to charge the affected tRNAs with the amino acid cognate to the ancestral tRNA, integrating the wrong amino acid during translation.

**Figure 6.**
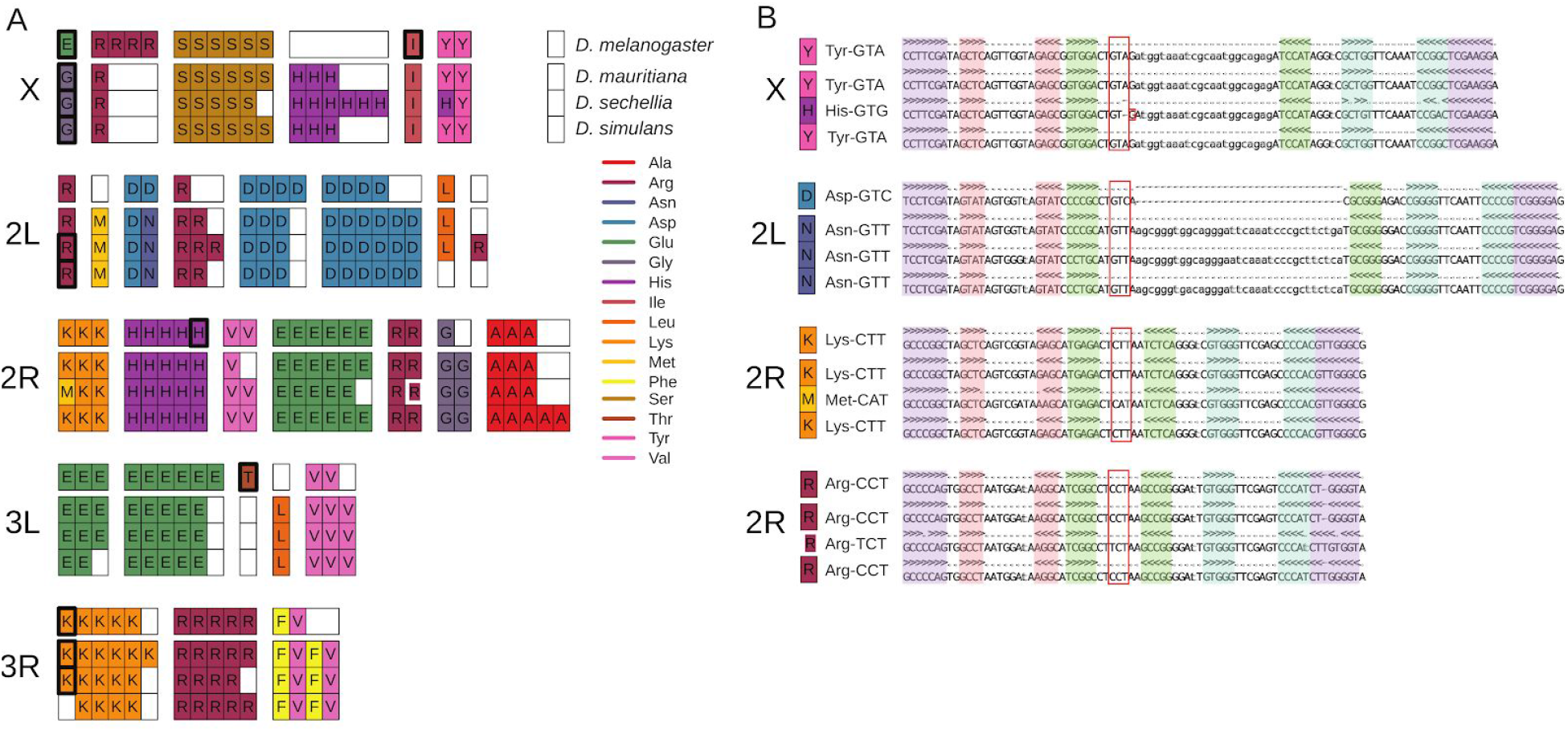
A) The subset of all nuclear tRNAs that differ in copy-number, isotype identity, or anticodon sequence between four mel-complex species. Each box represents an individual tRNA gene copy located within a larger syntenic cluster of tRNAs (grouped together as colored columns). Thick black outlines show tRNAs predicted to be pseudogenes. The thick white outline shows an arginine tRNA on Chromosome 2R predicted to use a different anticodon. B) Secondary structure alignments of orthologous nuclear tRNAs that exhibit anticodon shifts. The tRNA anticodon (red box), acceptor stem (purple), D arm (red), anticodon arm (green), and T arm (blue) are highlighted in the alignments. See Figure S24 for relative position on the chromosomes.

### Y Chromosome evolution

We identified Y-linked contigs in the sim-complex genomes using *D. melanogaster* Y-linked genes as queries. Y-linked contigs were short (<1 Mb) and lacked some homologous exons present in raw reads (e.g. exons 8–10 of *kl-3* and exons 6–8 of *kl-5*; supplemental Table S13; see also (Krsticevic et al. 2015; Chang and Larracuente 2019)), highlighting the challenges of assembling Y chromosomes even with long-read sequencing. We recovered 66, 58, and 64 of 83 *D. melanogaster* Y-linked exons (70–80%; supplemental Table S13) in *D. mauritiana, D. simulans*, and *D. sechellia*, respectively. A previous study found a duplication involving the Y-linked *kl-2* gene in *D. simulans (Kopp et al. 2006)*. We find that all known Y-linked genes, except *Ppr-Y*, exist in multiple copies in at least one of the sim-complex assemblies, and one exon of *Ppr-Y* appears duplicated in *D. mauritiana* raw long reads. Most duplication events correspond to partial tandem duplications (all but *ARY, Pp1-Y1, and Pp1-Y2*). We validated one duplicated exon from each of 10 Y-linked genes using PCR re-sequencing (except *Pp1-Y1*, which lacked mutations differentiating copies) (supplemental Table S13–14). Some duplicated exons (e.g. *kl-5* exons 9 and 10) are shared among sim-complex species, while other exons vary in copy number among species. For example, *ARY* is single-copy in *D. melanogaster* and *D. simulans*, but present in >3 copies in *D. sechellia* and *D. mauritiana*.

We identified 41 duplications from other chromosomes to the Y Chromosome only in the sim-complex species (supplemental Table S15), including 30 duplications not previously identified (Tobler et al. 2017). Among the 41 Y-linked duplications, 22 are shared by at least 2 sim-complex species and likely originated in the ancestor of the sim-complex. We verified putative Y-linked duplicates with PCR, confirming male-specificity for 16 of 17 of tested duplications (supplemental Table S14–15). We found that the Y Chromosomes of sim-complex species share an insertion derived from mtDNA that is absent in *D. melanogaster*.

## DISCUSSION

Here we uncover novel structural variation in both euchromatin and highly repetitive pericentromeric regions of the *D. simulans* species complex. This variation is substantial: approximately 15% of sim-complex genomes are not 1:1 orthologous with *D. melanogaster*, more than twice the number of nucleotide substitutions between these genomes (Begun et al. 2007). We find most rearrangements in heterochromatic genomic regions (Jagannathan et al. 2017; Sproul et al. 2020) likely influenced by both the density of repetitive DNA and the scarcity of genes. The former renders DNA repair mechanisms mutagenic, creating rearrangements, while the latter reduces selection against rearrangements in these regions. Such heterochromatic rearrangements may play a role in speciation, as many factors linked to genetic incompatibilities between species are located in pericentromeric heterochromatin (Bayes and Malik 2009; Cattani and Presgraves 2009; Ferree and Barbash 2009)

We also discovered 62 tandem duplications present only in the sim-complex genomes that duplicate one or more protein-coding exons. Such mutations frequently contribute to adaptation, functional innovation, and genetic incompatibilities (Lynch and Force 2000; Long et al. 2003; Ting et al. 2004; Arguello et al. 2006; Katju and Lynch 2006; Tao et al. 2007b; Zhou et al. 2008; Chakraborty and Fry 2015; Helleu et al. 2016; Eickbush et al. 2019). In the branch leading to the sim-complex, the rate of new gene acquisition is roughly one new gene every 78,000 years for full duplicates (∼12.8 duplicates per my) or one new gene per 40,000 years for partial gene duplicates (∼24.8 duplicates per my). The lower bound of these rates (1-2×10^−9^ new genes / gene / year) is consistent with previous estimates over a different timescale (Osada and Innan 2008). These estimates suggest that the rate of new gene acquisition per single copy gene is similar to the per nucleotide neutral mutation rate (Keightley et al. 2014). The proportion of exonic duplicates fixed in the sim-complex branch is greater than the proportion of polymorphic exonic duplicates in *D. simulans* (0.72 vs 0.408, proportion test, p-value = 3.41×10^−9^)(Rogers et al. 2014), whereas the proportion of intergenic (i.e. putatively nonfunctional) duplicates shows the opposite pattern (0.28 vs 0.43, proportion test, p-value = 0.0029). This suggests that either the exonic duplicates accumulated under positive selection in the sim-complex ancestral lineage or the polymorphism data, which is based on short reads, is missing duplicates. Further study with polymorphism data from highly contiguous *D. simulans* genome assemblies will resolve this puzzle.

These *Drosophila* genomes differ in TE content and composition, likely due to historical and ongoing differences in TE activity, natural selection, and host genome repression. Approximately 75–80% of TE content in all four genomes is due to species-specific insertions (Fig. 4), which are likely polymorphic within species (Chakraborty et al. 2018). This is consistent with most TE content resulting from recent activity (*Drosophila* 12 Genomes Consortium 2007; Lerat et al. 2011; Kofler et al. 2015; Bargues and Lerat 2017). Non-LTR retrotransposons comprise the majority (52%) of the old TEs found in all four mel-complex species, whereas DNA transposons comprise most (71%) of the younger fixed TE sequences found only in the sim-complex species. The widespread INE-1 DNA element (Quesneville et al. 2005; Yang and Barbash 2008; Sackton et al. 2009) is far more prevalent in the sim-complex ancestor than in the mel-complex ancestor, suggesting a burst of INE-1 activity in the sim-complex after diverging from *D. melanogaster*. On the other hand, *D. melanogaster’*s genome is enriched for LTR elements due to recent TE activity in this lineage (Bowen and McDonald 2001; Bergman and Bensasson 2007; Kofler et al. 2015). These LTRs have increased the size of *D. melanogaster’s* genome through both intergenic and intragenic insertions, so that euchromatic introns containing repetitive DNA are ∼10% longer in *D. melanogaster* than sim-complex species (supplementary text). However, while the sim-complex does harbor less TE content than *D. melanogaster* (Fig. 4A) (*Drosophila* 12 Genomes Consortium 2007), we observe only ∼17% less total TE sequence in *D. simulans* than in *D. melanogaster*, which is substantially lower than the consensus (Young and Schwartz 1981; Dowsett and Young 1982; Nuzhdin 1995; Vieira et al. 1999; Vieira and Biémont 2004; *Drosophila* 12 Genomes Consortium 2007).

Intron size evolution may also be modulated by differences in insertion and deletion mutations (Petrov et al. 1996; Petrov and Hartl 1998; Blumenstiel et al. 2002), recombination rates (True et al. 1996; Brand et al. 2018), effective population sizes (Kofler et al. 2012), or variation in constraint mediated by the presence of conserved noncoding elements (Manee et al. 2018). Further study is needed to determine which factors contribute to the differences between sim-complex genomes; for example, among the sim-complex species, *D. sechellia* has the longest complex introns in heterochromatin, but not in euchromatin (Supplemental Table S10), which could be a result of both low recombination rates in heterochromatin and the small effective population size of this species (Kliman et al. 2000; McBride 2007; Singh et al. 2007). A small effective population size in *D. sechellia* might also lead to the enrichment of tRNA anticodon shifts (75% of all observed) and expansion of euchromatic satellites.

TE activity is deleterious (Petrov et al. 2011; Cridland et al. 2013; Chakraborty et al. 2019): transposition disrupts genes and other functional elements (e.g. supplemental Fig. S25) (Cooley et al. 1988), TE sequences can act as ectopic regulatory elements (Feschotte 2008), and provide templates for ectopic recombination (Montgomery et al. 1987; Miyashita and Langley 1988). Like other eukaryotes, *Drosophila* has evolved host defenses against TE proliferation (Aravin et al. 2007; Brennecke et al. 2007; Chung et al. 2008; Kelleher et al. 2018). Interspecific differences in these host defenses may contribute to the TE abundance differences between the sim-complex and *D. melanogaster*. TE insertions also alter local chromatin state in *Drosophila*, which can spread and suppress the expression of adjacent genes, with potentially deleterious consequences (Lee and Karpen 2017). Heterochromatin proteins are expressed at higher levels in *D. simulans* than *D. melanogaster*, which may cause heterochromatin to spread further from TEs into nearby regions in *D. simulans* (Lee and Karpen 2017). Thus, selection to eliminate euchromatic TE insertions may be stronger in *D. simulans* than in *D. melanogaster*, contributing to the excess of TEs in the latter. We identified a recent duplication of *su*(*f*)–a suppressor of *Gypsy* LTR retrotransposon expression–in all sim-complex species (Parkhurst and Corces 1986; Mazo et al. 1989). The extra copy could contribute to the lower activity and prevalence of LTR elements in the sim-complex species compared to *D. melanogaster* (Fig. 4).

Sex chromosomes play a special role in the evolution of postzygotic hybrid incompatibilities (Coyne and Orr 1989). We find that euchromatic duplications, deletions, and inversions are enriched on the X Chromosome (supplemental Table S11): 90% of all rearrangements between sim-complex genomes are X-linked (supplemental Table S8). We also report an enrichment (∼15-to-50-fold) of X-linked satellite sequences, exceeding even previous reports (∼7.5-fold; (Garrigan et al. 2014)). Ectopic exchange between repeats during DNA repair can create genomic rearrangements. X-linked euchromatic satellites may contribute to the enrichment of rearrangements on this chromosome (Fig. 2A and Supplemental Table S9; (Sproul et al. 2020)). It remains unclear whether these rearrangements contribute to the enrichment of hybrid incompatibility factors on the X Chromosomes within the sim-complex (Tao and Hartl 2003; Masly and Presgraves 2007). The sim-complex genomes also contain a duplication of *mh*, whose protein product interacts with the X-linked heterochromatic satellite called *359-bp*–a member of the *1*.*688 gm/cm*^*3*^ satellites–to maintain genome stability during embryogenesis (Loppin et al. 2001; Tang et al. 2017; Delabaere et al. 2014). The derived copy of *mh* produces a shorter transcript than the ancestral copy, has male-biased expression, and likely binds to *359-bp*, given the similarity between the ancestral and derived proteins (supplemental Fig. S21). We speculate that the duplicated *mh* may play a role in the male germline regulating *359-bp* related satellites that have proliferated across the sim-complex species X Chromosomes (Jagannathan et al. 2017; Sproul et al. 2020).

Despite harboring few genes, the *Drosophila* Y Chromosome contributes to hybrid incompatibilities and affects phenotypes including longevity, immunity (Araripe et al. 2016; Case et al. 2015; Kutch and Fedorka 2015; Brown et al. 2020), meiotic drive (Voelker 1972; Atlan et al. 1997; Unckless et al. 2015), male fitness (Chippindale and Rice 2001) and gene expression across the genome (Lemos et al. 2010; Branco et al. 2013). We discovered extensive divergence between mel-complex species in the genic content of Y chromosomes resulting from rampant inter- and intrachromosomal duplication. Y-linked gene content in *Drosophila* is shaped by gene duplication from the autosomes (Kopp et al. 2006; Koerich et al. 2008; Carvalho et al. 2015; Ellison and Bachtrog 2019). We detect 41 duplications from the other chromosomes to sim-complex Y chromosomes. We also discovered that nearly all Y-linked genes are duplicated in at least one species. This amplification of Y-linked genes appears to be a common feature of *Drosophila* Y Chromosomes, and may reflect a strategy to compensate for the heterochromatic environment or ongoing genetic conflict with the X Chromosome (Kopp et al. 2006; Koerich et al. 2008; Carvalho et al. 2015; Ellison and Bachtrog 2019).

The structural divergence between these species extends to the endosymbionts they carry. We uncovered extensive structural evolution in *Wolbachia* genomes between *w*Mau and the corresponding *D. simulans Wolbachia* strains (supplementary Fig. S8A–C). Further study is necessary to understand whether such variants affect important phenotypes like titer and transmission (Meany et al. 2019, Serbus and Sullivan 2007), virulence (Chrostek and Teixeira 2018), fitness (Turelli and Hoffmann 1995; Kriesner et al. 2013), *Wolbachia* frequency variation (Cooper et al. 2017, Kriesner et al. 2016), or cytoplasmic incompatibility (Hoffmann and Turelli 1997).

Previous assemblies were biased towards unique sequences, neglecting repetitive regions (*Drosophila* 12 Genomes Consortium 2007; Bhutkar et al. 2008; Garrigan et al. 2012; Hu et al. 2013). However, these regions harbor extensive hidden genetic variation relevant to genome evolution and organismal phenotypes (Khost et al. 2017; Chakraborty et al. 2018; Stein et al. 2018; Chaisson et al. 2019; Chang and Larracuente 2019; Stitzer et al. 2019; Chakraborty et al. 2020; Miga et al. 2020). Understanding the evolution of these rapidly diverging repetitive, complex genomic regions and their effects on adaptation and species differentiation requires a direct comparison between closely related species. Here we show that the genomes of these four *Drosophila* species have diverged substantially in the regions that have been previously recalcitrant to assembly. Future studies of interspecific variation in genome structure will shed light on the dynamics of genome evolution underlying speciation and species diversification.

## METHODS

### Data collection

Unless otherwise stated, we use the following strains: *D. mauritiana* (*w*12), *D. simulans* (*w*^XD1^), and *D. sechellia* (Rob3c / Tucson 14021-0248.25) (Garrigan et al. 2012; Meiklejohn et al. 2018). We extracted gDNA following Chakraborty et al. (2016). The standard 20-kb library protocol was carried out at the UCI genomics core using the P6-C4 chemistry on PacBio RS II.

To collect RNA sequencing, flies from the sim-complex species were reared at room temperature on standard cornmeal-molasses medium. We collected 20-30 3-5 days old virgin males and females, and dissected testes from at least 100 males. For *D. simulans* and *D. mauritiana*, total RNA was extracted using TRIzol (Invitrogen) and phase-lock gel tubes (Fisher Scientific). Sequencing libraries generated by Illumina TruSeq Stranded mRNA kit were sequenced at the University of Minnesota Genomics Center. For *D. sechellia*, we isolated total RNA using the RNeasy Plus Kit (Qiagen), and constructed libraries using TruSeq RNA Sample Preparation Kit V2 (Illumina) with oligo-dT selection (data available in PRJNA541548).

### Genome assembly

#### Nuclear genome assembly

We assembled the nuclear genomes of the sim-complex species *de novo* following the previously described approaches for assembly and polishing (Supplemental Fig. S26)(Chakraborty et al. 2016). To detect possible misassemblies, we identified orthologs of all *D. melanogaster* heterochromatic genes using BLAST and examined their gene structure. Because there are no detectable interchromosomal rearrangements in the mel-complex species (Bhutkar et al. 2008), we flagged contigs with genes that translocated between chromosome arms or that appeared on more than two contigs, as potential misassemblies. We then manually fixed 10 misassemblies based on these results (supplemental Table S16; supplemental Fig. S27).

#### Mitochondrial genome assembly

We extracted raw reads mapping to an existing partial mitochondrial genome using BLASR (Chaisson and Tesler 2012). (https://github.com/mahulchak/mito-finder). We selected the longest read exceeding a length cutoff of 18 kb (the mitochondrial genome is approximately 19 kb) and trimmed the redundant sequences resulting from multiple polymerase passes through the SMRTbell template. Trimmed reads were polished twice with Quiver (Chin et al. 2013) to generate a consensus of all mitochondrial reads.

#### *Wolbachia* genome assembly

We took advantage of the fact that endosymbionts are co-sequenced with their hosts in shotgun sequencing data to assemble complete *Wolbachia* genomes from our PacBio data (Faddeeva-Vakhrusheva et al. 2017; Basting and Bergman 2019; Kampfraath et al. 2019). We identified a complete *Wolbachia* genome in *D. mauritiana* from the Canu assembly. For *D. sechellia*, we collected all reads mapping to two reference *Wolbachia* genomes (CP003884.1 and CP003883) using BLASR v5.1 (Chaisson and Tesler 2012) with parameters (--clipping soft --bestn 1 --minPctIdentity 0.70). We assembled these reads using Canu v1.3 with the parameters (genomeSize=3m; (Koren et al. 2017)). No *D. simulans* reads were mapped to the *Wolbachia* genomes.

### Assembly validation and quality control

We evaluated long read coverage to identify assembly errors and validate copy number variants. We mapped raw long reads to assemblies using *BLASR* (version 1.3.1.142244; parameters: -bestn 1 -sam; (Chaisson and Tesler 2012)) or minimap2 (2-2.8 parameters: -ax map-pb; (Li 2016)). We calculated long read coverage across the contigs using the SAMtools mpileup and depth (*-Q 10 -aa*) commands. To validate CNVs, we chose 20 random CNVs for each species and inspected long read coverage across the regions containing CNVs following (Chakraborty et al. 2018). The presence of at least 3 long reads spanning the entire CNV was classified as evidence supporting the variant.

We used the script in Masurca v3.2.1 (Zimin et al. 2013) to identify redundant sequences in our assemblies. We designated contigs as residual heterozygosity candidates (those greater than 40 kb require >90% identity and those between 10 and 40 kb require >95% identity to the longest contigs). To detect microbial contamination in our assemblies (supplemental Table S7), we used BLAST+ v2.6.0 (Altschul et al. 1990) with blobtools (0.9.19.4; (Laetsch and Blaxter 2017)) to search the NCBI nt database (parameters “ -task megablast -max_target_seqs 1 -max_hsps 1 -evalue 1e-25”) and calculated the Illumina coverage of all contigs for *D. mauritiana, D. simulans* and *D. sechellia*, respectively (supplemental Table S17; supplemental Fig. S28).

We applied the method of Koren et al. (Koren et al. 2018) to the polished, pre-scaffolded assemblies to estimate base level error rates from the concordance between Illumina reads and an assembly of the same strain (i.e. QV). We calculated BUSCOs in our assemblies with BUSCO v3.0.2 against the Diptera database (Waterhouse et al. 2017). Some duplicated BUSCOs in *D. simulans* remained due to persistent alternate haplotigs. We inspected these 71 duplicate BUSCOs, identifying 58 with one member on Muller element contigs and the others on smaller, putative alternate haplotigs. BUSCO metrics were re-calculated without these unplaced contigs (Table S5). We also applied QUAST v5.0.2 (Mikheenko et al. 2018) to evaluate the quality of assemblies based on the mapping status of Illumina data. For *D. simulans* and *D. sechellia*, we used independently-generated male and female reads (Wei et al. 2018) to avoid the ascertainment bias due to the Illumina reads used in polishing our assemblies (supplemental Table S17). For *D. mauritiana*, we used the female Illumina reads for both our assembly and the previous assemblies (Garrigan et al. 2014).

### Scaffolding

We scaffolded the assemblies with mscaffolder (https://github.com/mahulchak/mscaffolder) following (Chakraborty et al. 2018) using *D. melanogaster* as the reference. Scaffolded contigs were joined with 100 Ns and unscaffolded contigs were prefixed with ‘U’.

### Annotation

#### Transcript annotation

We mapped transcripts and translated sequences from *D. melanogaster* (r6.14) to each assembly using MAKER2 (v2.31.9; (Holt and Yandell 2011). We also generated RNA-seq from whole females, whole males, and testes from the sim-complex species. We mapped this data (see details in supplemental Table S18) using HISAT2.1.0 with the MAKER2 annotation, and then used StringTie 1.3.4d to generate consensus annotations (Pertea et al. 2016). We further annotated putative duplicated genes in *D. simulans* using Iso-Seq data from (Nouhaud 2018). We applied the IsoSeq3 pipeline (v3.1.2) to correct and polish the raw reads, then generated full length cDNA sequences (Gordon et al. 2015). Polished cDNA sequences were mapped to the assembly using minimap2 (r2.16, (Li 2016)) with parameters “ -t 24 -ax splice -uf --secondary=no -C5”. We then used cdna-cupcake (v10.0.1 with the parameter “ --dun-merge-5-shorter”; https://github.com/Magdoll/cDNA_Cupcake) to cluster the isoforms in the cDNA alignment and transfer it to the annotation. We used BLAST (-evalue 1e-10; (Altschul et al. 1990) homology to assign the predicted transcripts to *D. melanogaster* transcript sequences. To identify conserved introns, we kept isoforms with the same numbers of exons and only used introns flanked by exons of similar size (within 10% length difference) in each species. To compare intron sizes between species, we used the longest isoform from each gene. We also annotated 61 introns from 6 genes with large introns (> 8 kb) based on BLAST results.

#### Large structural variant detection

To identify large scale synteny, we created whole-genome alignments with the Mauve aligner (build 2015-2-13) using the progressiveMauve algorithm (Darling et al. 2010) with the default parameters: default seed weight, determine LCBs (minimum weight = default), full alignment with iterative refinement. We plotted gene density based on Dm6 annotations in *D. melanogaster* was plotted using Karyoploter (Gel and Serra 2017)).

#### Annotation of repetitive elements

We annotated new complex satellites using Tandem Repeat Finder and annotated novel TEs using the REPET TE annotation package (Flutre et al. 2011) (supplemental Fig. 29A, B). We removed complex satellite annotations from the *Drosophila* Repbase release (20150807), and combined the rest of the library with our newly annotated satellites and TEs. We then updated repeat classifications (supplemental Fig 29C) and used the resulting library (Supplemental File S1) to annotate the three sim-complex species and the *D. melanogaster* reference with RepeatMasker v4.0.5 (Smit et al. 2013)(supplemental Fig. 29).

We calculated the proportion of each repeat family and the proportion of TEs that are DNA transposons, non-LTR, and LTR retrotransposons in 100-kb windows across the scaffolds containing major chromosome arms. We determined approximate euchromatin/heterochromatin boundaries in the major scaffolds based on boundaries from *D. melanogaster* (Hoskins et al 2015) in each sim-complex assembly. We considered Chromosome 4 and all unassigned contigs to be heterochromatin. TE sequence annotations in our *D. simulans* assembly were called exonic when they fell inside the alignment between the Iso-Seq transcript and the genome.

#### tRNA annotation and analysis

We used tRNAscan-SE v1.4 (options: -H; (Lowe and Eddy 1997) to annotate tRNAs and predict secondary structures in the *D. melanogaster* reference (r6.09) and in sim-complex assemblies. We sorted tRNAs by position and represented them as peptide sequences based on the predicted tRNA isotype that we aligned using MUSCLE v3.8.31 (Edgar 2004). We inspected these coarse alignments of tRNA positions for each chromosome (X, 2L, 2R, 3L, 3R) using conservation of gene order, strand orientation, inter-tRNAs distances, anticodon sequence, and intron positions to identify positional tRNA orthologs within syntenic clusters (see supplementary text). We also used a BLAST-based orthology discovery method—similar to methods described in Rogers et al (2010)—to map tRNAs from *D. mauritiana, D. sechellia*, or *D. simulans* that did not share positional orthologs with tRNAs in *D. melanogaster* (see supplementary text).

#### Genomewide SV annotation

We aligned each member of the sim-complex to *D. melanogaster* (Hoskins et al. 2015) using MUMmer 4.0 (NUCmer -maxmatch) (Marçais et al. 2018) and LASTZ (Harris 2007). MUMmer alignments were processed using *SVMU* v0.3 (Structural Variants from MUMmer; https://github.com/mahulchak/svmu commit 9a20a2d; (Chakraborty et al. 2018, 2019) to annotate the SVs as duplicates originating in either the sim-complex or in *D. melanogaster*. We added duplications that MUMmer failed to recover using an approach based on LASTZ alignments (Schwartz et al. 2003) and UCSC Genome Browser alignment chaining. The LASTZ/axtChain workflow is available at https://github.com/yiliao1022/LASTZ_SV_pipeline (Kent et al. 2003). Additional details are provided in the supplement (see supplementary section “ SV Validation and analysis”).

### Shared TE analysis

We limited the shared TE analysis to euchromatic regions. To identify TEs shared between species, we performed all pairwise alignments of the sim-complex species to each other and to *D. melanogaster* using NUCmer -maxmatch -g 1000 in MUMmer v4. We extracted syntenic regions from alignment with svmu 0.3 and validated these regions by inspecting the dotplots (supplemental Fig. S30). To identify TE sequences completely contained with syntenic regions between species pairs, we used BEDTools (BEDtools -u -f 1.0 -a te.bed -b cm.eu.txt) (Quinlan and Hall 2010). We identified TEs shared among all four mel-complex species using the D. *mauritiana* genome as the reference. TEs shared between *D. mauritiana–D. sechellia* (A) and *D. mauritiana*–*D. simulans* species pairs (B) were inferred to be derived from either the sim-complex or mel-complex ancestral lineages (Fig. 4), whereas TEs shared between A, B, and *D. mauritiana*–*D. melanogaster* pair were inferred to be derived from the TEs fixed only in the mel-complex ancestral lineage (BEDtools intersect -u -a te.simclade.bed -b te.dmau-dmel.bed). We report differences in the abundance of existing TE families within these genomes and make no inferences that TEs are restricted to or missing from any subset of these four species.

### Y Chromosome analyses

We used BLAST to identify the orthologs of all known *D. melanogaster* Y-linked genes in the sim-complex assemblies (Altschul et al. 1990). The sequences of new Y-linked genes were extracted based on BLAST results. We inspected all alignments of duplicates to ensure that Y-linked duplicates are distinct from the parental copies.

### Cytological validation

We conducted FISH following the protocol from (Larracuente and Ferree 2015). Briefly, brains from third instar larva were dissected and collected in 1X PBS, followed by an 8-min treat of hypotonic solution (0.5% sodium citrate), fixed in 1.8% paraformaldehyde, 45% acetic acid, and dehydrated in ethanol. The 193XP probe was made by IDT with 5’-/56-FAM/ACATTGGTCAAATGTCAATATGTGGTTATGAATCC-3’ (supplemental Table S14). Slides are mounted in Diamond Antifade Mountant with DAPI (Invitrogen) and visualized on a Leica DM5500 upright fluorescence microscope, imaged with a Hamamatsu Orca R2 CCD camera and analyzed using Leica’s LAX software.

## DATA ACCESS

All raw genomic data and RNA-seq have been deposited to NCBI. The accession numbers of the assemblies, Illumina and Pacific Biosciences raw reads are provided in supplemental Table S17.

## DISCLOSURE DECLARATION

The authors do not declare any conflict of interest.

## ACKNOWLEDGMENTS

This work was funded by the National Institutes of Health (K99GM129411 to M.C., R35GM119515 to A.M.L., R01GM123303 to J.J.E., R01GM123194 to CDM) and National Science Foundation (NSF MCB 1844693 to A.M.L., NSF DDIG 1209536 to J.V., IOS-1656260 to J.J.E., NSF GRFP 1342962 to J.R.A.) and funding from the University of Nebraska-Lincoln to C.D.M and K.L.M. and the University of California, Irvine to J.J.E. A.M.L. was supported by a Stephen Biggar and Elisabeth Asaro fellowship in Data Science. C.-H.C. was supported by the Messersmith Fellowship from the University of Rochester and the Government Scholarship to Study Abroad from Taiwan. This work was made possible, in part, through access to the Genomics High-Throughput Facility Shared Resource of the Cancer Center Support Grant CA-62203 at the University of California, Irvine, and NIH shared-instrumentation grants 1S10RR025496-01, 1S10OD010794-01, and 1S10OD021718-01. We also thank the University of Rochester Center for Integrated Research Computing for access to computing cluster resources. We would like to thank Drs. Daniel Garrigan and Sarah Kingan for generating *D. sechellia* transcriptome data. We would also like to thank Nishant Nirale, Luna Thanh Ngo, and Cécile Courret for help with data collection and management, and Brandon Cooper, Christina Muirhead, Robert Kofler, Grace Lee, and Casey Bergman for comments on the manuscript.

## Notes

### Competing Interest Statement

The authors have declared no competing interest.

### Summary of Updates

The new version has extensive revisions of the text and contains revised figures.

