## Supplementary information for "Evolution of genome structure in the *Drosophila simulans* species complex"

### Supplementary methods and analyses

#### Validating Assemblies by BAC sequences

To validate our assemblies, we mapped BAC sequences (DSM1 and DSE1 from (Murakami et al. 2008)) to our assemblies of *D. simulans* and *D. sechellia*, respectively. In *D. simulans*, 6673 of 6958 BACs (95.9%) are mapped to a region < 300 kb (supplemental Fig. S6A). We also detect that in 43 (0.6%) BACs, the sequences from two ends are mapped to two different chromosome arms or do not map next to each other (> 300kb). In *D. sechellia*, 6433 of 6550 BACs (98.2%) are mapped to a region < 300 kb (supplemental Fig. S6B). We also detect that in 35 (0.5%) BACs, the sequences from two ends are mapped to two different chromosome arms or do not map next to each other (> 300kb). Our assemblies are consistent with the BACs while the strains used for BAC libraries are different from our sequenced strains.

#### tRNA annotation and analysis

Using our manually curated alignments of tRNAs predicted by tRNAscan-SE v1.4 (options: -H; (Lowe and Eddy 1997), we identified syntenic blocks of neighboring tRNAs separated by either conserved (*i.e.*, present in all species) tRNAs of a different isotype or by large physical distances along the chromosome. From these syntenic blocks, we identified changes in copy-number, isotype identity, anticodon sequence, or pseudogene designation (as predicted by tRNAscan-SE). We refer to the tRNAs within these syntenic blocks as positional orthologs, though we caution that many of these tRNAs may have arisen through duplications or more complicated local rearrangements and 1:1 orthology between any two tRNA positional orthologs is not implied. We used raw long reads to verify nucleotide changes in predicted tRNA isotypes or anticodons within syntenic blocks, as some of these changes were the result of single-base substitutions or small indels in the tRNA anticodon loop and may be highly sensitive to errors in sequencing or mapping. Visualization of the alignments of raw reads at this position using the Integrative Genome Viewer or IGV (Robinson et al. 2011) revealed that none of the observed changes in tRNA isotypes or anticodons among our assembled genomes were the result of sequencing or mapping error.

For tRNAs from sim-complex species that did not share positional orthologs with tRNAs in *D. melanogaster*, we used a BLAST-based orthology discovery method—similar to methods described in Rogers et al (2010) Specifically, we asked if sequences flanking these tRNAs had orthologous sequences in *D. melanogaster* and if these sequences overlapped annotated tRNA genes in *D. melanogaster*. We first masked tRNA positions in each query assembly of sim-complex species using the maskfasta function in BEDtools v2.20.1 (default options) (Quinlan and Hall 2010). We then masked repetitive sequences in the *D. melanogaster* reference using RepeatMasker v4.0.5 (Smit et al. 2013) (options: -species drosophila -no_is)—which served as our custom BLAST database. We extracted a 10-kb region of sequence—5-kb from each flank (including neighboring masked tRNAs)—surrounding each tRNA of interest and searched against the repeat-masked *D. melanogaster* database using BLASTN v.2.2.29 (options: -max_hsps 10000 -evalue $10^{-10}$). Orthologous windows were identified when both the left- and right-flanking query sequences produced significant search hits separated by fewer than 20-Kbp in *D. melanogaster*. Putatively orthologous tRNAs were then identified if these orthologous windows either overlapped or flanked a tRNA annotated in *D. melanogaster*.

#### SV annotation, validation and analysis

We tested the enrichment of the duplicates overlapping full genes on the X chromosome using the exact binomial test in R. We calculated the proportion of total genes on the euchromatic X chromosome (2244/13861) based on the *D. melanogaster* GFF file (r6.09). We required a mutual 50% overlap between sim-complex species and *D. yakuba* to classify SVs as orthologs (BEDtools intersect -f 0.5 -F 0.5). We annotated inversions and insertions (>100 bp) using SVMU. The sim-complex species genomes were aligned to the *D. melanogaster* ISO1 reference genome using MUMmer (v4) and the alignment delta file for each sim-complex species was processed with SVMU (e.g. SVMU sim2mel.delta mel.fasta sim.fasta 100 l). The inversion breakpoints were extracted from the sv.txt output file using the 'INV' tag and the insertion coordinates were extracted using the 'INS' tag. To complement inversion detection by the automated pipeline, we also extracted all inverted mummer alignments (i.e. query start is bigger than query end) from the 'cords.txt' output from SVMU. The "cords.txt" file reports all mummer alignments from a delta alignment file in a tab-separated format. We used BEDtools to extract inverted query intervals that were not found by SVMU (BEDtools intersect -v -a output_from_cords.txt -b output_from_svmu). We filtered out inverted intervals that are TEs using BEDtools (BEDtools intersect -v -F 0.95 -a inversion_candidates.txt -b te.txt). This final TE filtered set of inversion candidates were further visually inspected in the whole-genome-alignment dotplots and was used for downstream analysis. To identify species-specific TE insertions in our reference strains, we identified TE insertions in SVs with at least 80% of full TE length (BEDtools intersect -u -f 0.8 -a ins.A.bed -b ins.B.bed). To infer the lineages where the inversions occurred, we examined the presence of the inversions in the FlyBase assemblies of the outgroup species *D.*

*yakuba* and *D. erecta*. To validate these inferences, we further aligned the *D. melanogaster* assembly to the FlyBase assembly of *D. annanassae* by their protein sequences (promer mel.fasta ana.fasta) and inspected the arrangement of protein-coding genes at the inversion breakpoints. To identify the duplication CNVs, we employed both MUMmer (SVMU) and lastz (https://github.com/yiliao1022/LASTZ_SV_pipeline) based pipelines and combined their results using bedtools. In particular, we used the 'CNV-Q' tags in SVMU outputs to pull out the >100 bp *D. melanogaster* sequences that were duplicated in the sim-complex species. To combine the LASTZ and SVMU duplication output, we identified the SVMU-specific calls (BEDtools intersect -v -a svmu_calls.txt -b lastz_calls.txt) and added them to the LASTZ output. We used both aligners because LASTZ is more sensitive at aligning diverged sequences whereas MUMmer excels in less divergent contexts. To validate the duplicates, we aligned the long reads to the assemblies using BLASR (Chaisson and Tesler 2012) and inspected the alignment bam files in IGV (Thorvaldsdóttir et al. 2013). Presence of at least 3 reads spanning an entire duplication was used as the evidence to confirm the duplicate.

#### Mutation pattern analyses

Introns can harbor functional elements like splice signals which determine a minimum size for introns, or regulatory sequences, which are common in larger introns (Chorev and Carmel 2012). Consistent with these functions, introns tend to evolve under stronger purifying selection than synonymous sites (Bergman and Kreitman 2001; Haddrill et al. 2005; Andolfatto 2005). Local recombination rates may also affect intron size: the negative correlation between intron size and recombination rate suggests that large introns are weakly deleterious (Carvalho and Clark 1999), or alternatively, that they may be favored because they can decrease linkage between exons (Comeron and Kreitman 2000). Previous studies indicate that *D. simulans* introns are shorter than *D. melanogaster*'s (Comeron and Kreitman 2000; Ometto et al. 2005) perhaps in part because insertions in introns are favored in *D. melanogaster* (Presgraves 2006). Here we present a comprehensive study of intron length evolution in the four mel-complex species including heterochromatic introns. For our analysis, we considered 6,289 orthologous genes and analyzed 21,860 introns with conserved annotation positions in *D. simulans*, *D. mauritiana* and *D. sechellia*. We found that introns are significantly shorter in the sim-complex compared to *D. melanogaster* (paired t-test, $P < 2e\text{-}4$). However, we did not detect a length difference within the sim-complex (paired t-test, $P > 0.06$). We first examined the effect of indel mutation bias between *D. melanogaster* and *D. simulans* using population polymorphism data from highly inbred lines from the natural populations (10,462 polymorphic indels from *D. melanogaster* (DGRP; (Huang et al. 2014)) and 143,139 polymorphic indels from *D. simulans* (Signor et al. 2018)). Consistent with these studies, we find that both *D. simulans* and *D. melanogaster* introns have more segregating deletions than insertions (1.35 and 1.41 deletions per insertion, respectively). In *D. simulans,* deletions are on average larger than insertions (4.45bp vs. 3.31bp), while segregating insertions are slightly larger than deletions (7.19bp vs. 6.68bp) in *D. melanogaster*. These biases may explain why simple introns are ~3 bp longer in *D. melanogaster* genome than *D. simulans*. The similarity in simple intron size within the sim-complex species suggests that they share similar insertion and deletion biases.

#### Length of collinear groups

We extracted all 1:1 collinear groups across all mel-complex species from Mauve's results. We found that the length collinear groups are shorter in heterochromatin compared to euchromatin. This pattern is consistent with more genomic rearrangements happening in heterochromatin than euchromatin in the evolutionary history. The length of collinear groups is significantly shorter in heterochromatin than euchromatin for all autosomal arms (two-tail t tests, all $P < 1e-8$; Supplemental Fig. S9), but not for the X chromosome (two-tail t tests, all $P > 0.18$; Supplemental Fig. S9). The difference between the X chromosome and autosomes is due to the massive rearrangements on X chromosome euchromatin.

#### Estimation of orthologous genes in assemblies

We used best reciprocal BLASTx and tBLASTn hits with an e-value threshold 1e-10 to identify *D. melanogaster* orthologous genes in our assemblies. For identifying in *D. melanogaster* orthologs in previous assemblies, we applied the best reciprocal BLASTp to hits with the same e-value threshold 1e-10. We also include the annotation from (Rogers et al. 2014), and the result is shown in Supplemental Table S6.

### Supplementary figures

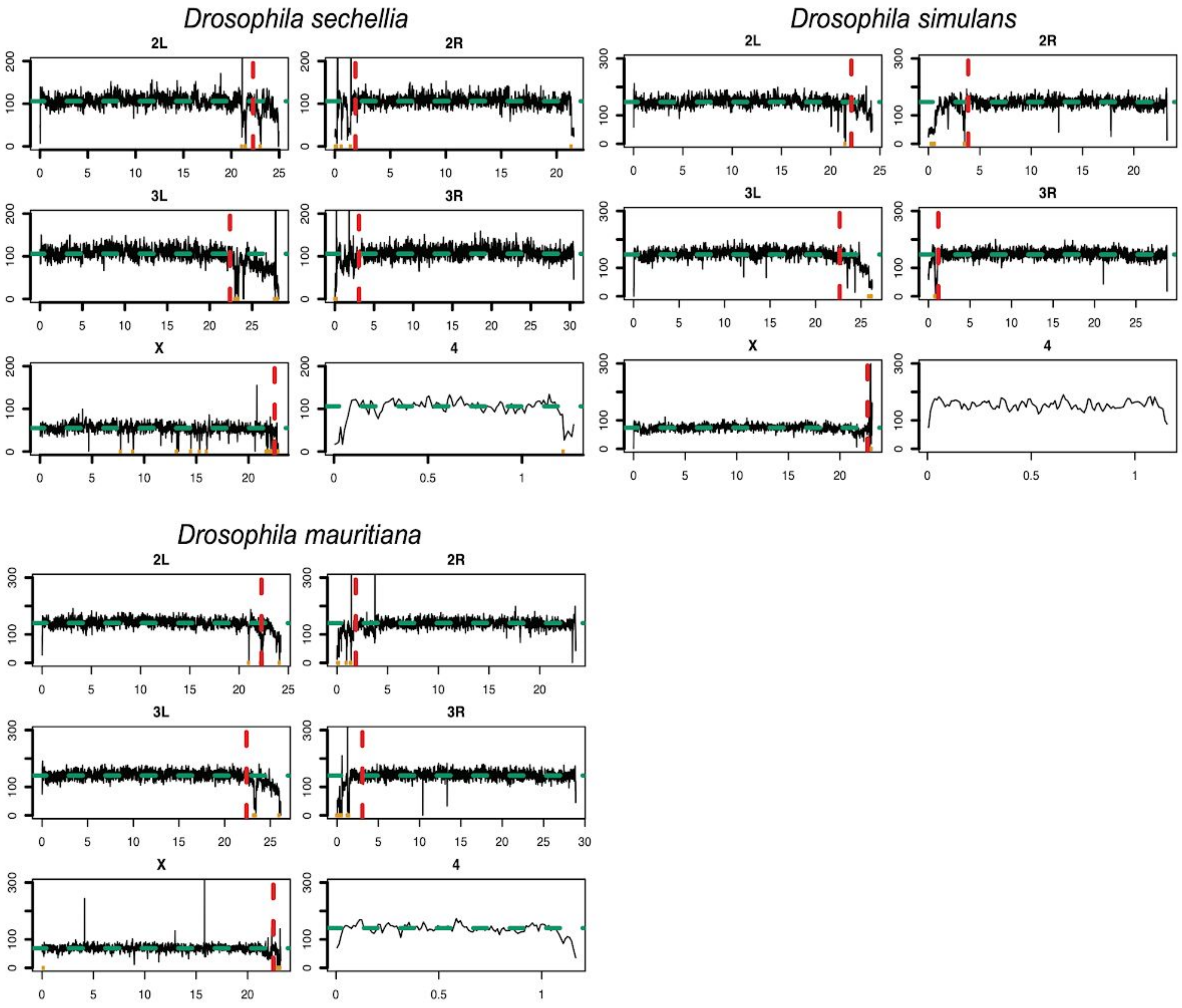


Figure S1. The median coverage of PacBio uniquely mapped reads (mapQ > 10) in every 10kb window across the assemblies in the sim-complex. The dotted green lines are the median coverage of autosomes and X chromosomes and the dotted redlines are the boundaries between euchromatin and heterochromatin. The orange dots are the gap position in the scaffolds.

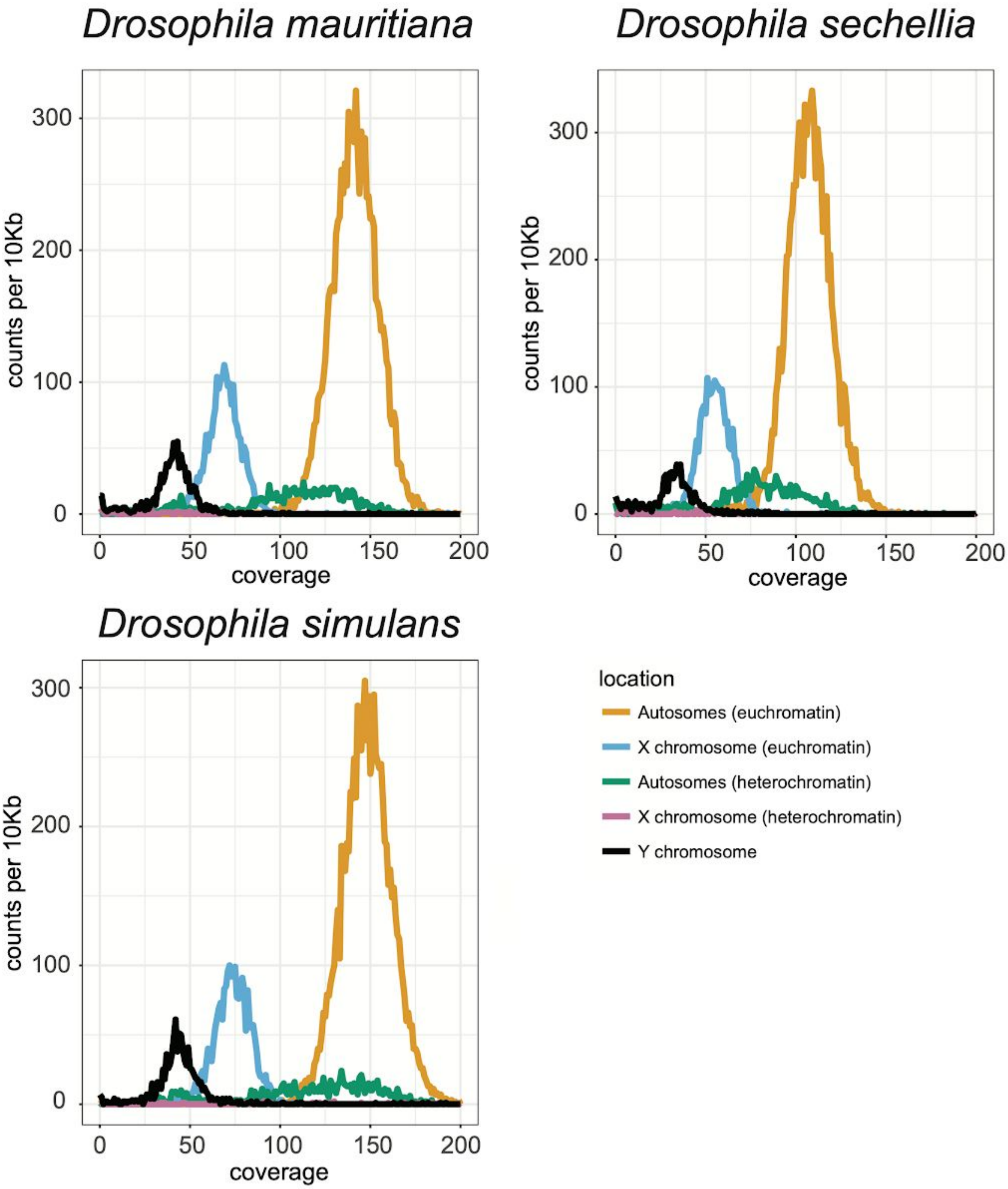


Figure S2. Histogram of coverage per 10kb window in the sim-complex. The different colors represent different regions of genomes (A: autosomes, X: X Chromosome, and Y: Y Chromosome). We isolated the heterochromatic regions (Ahet and Xhet) from euchromatic regions (A and X) to infer the sequencing bias.

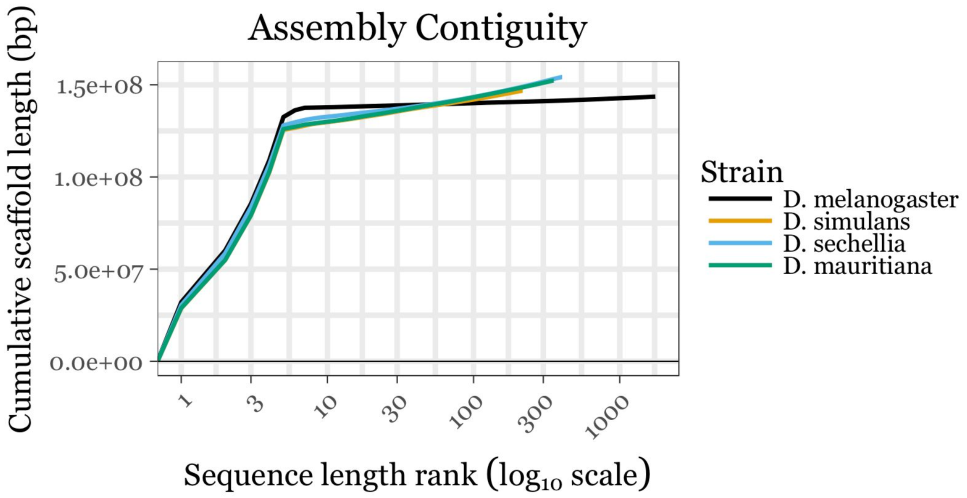


Figure S3. The contiguity of scaffolds in the assemblies from sim-complex species and *D. melanogaster* (R6). We ranked the scaffolds by their length and plotted their cumulative length from the longest contigs/scaffold to the shortest one. The different colors represent different species.

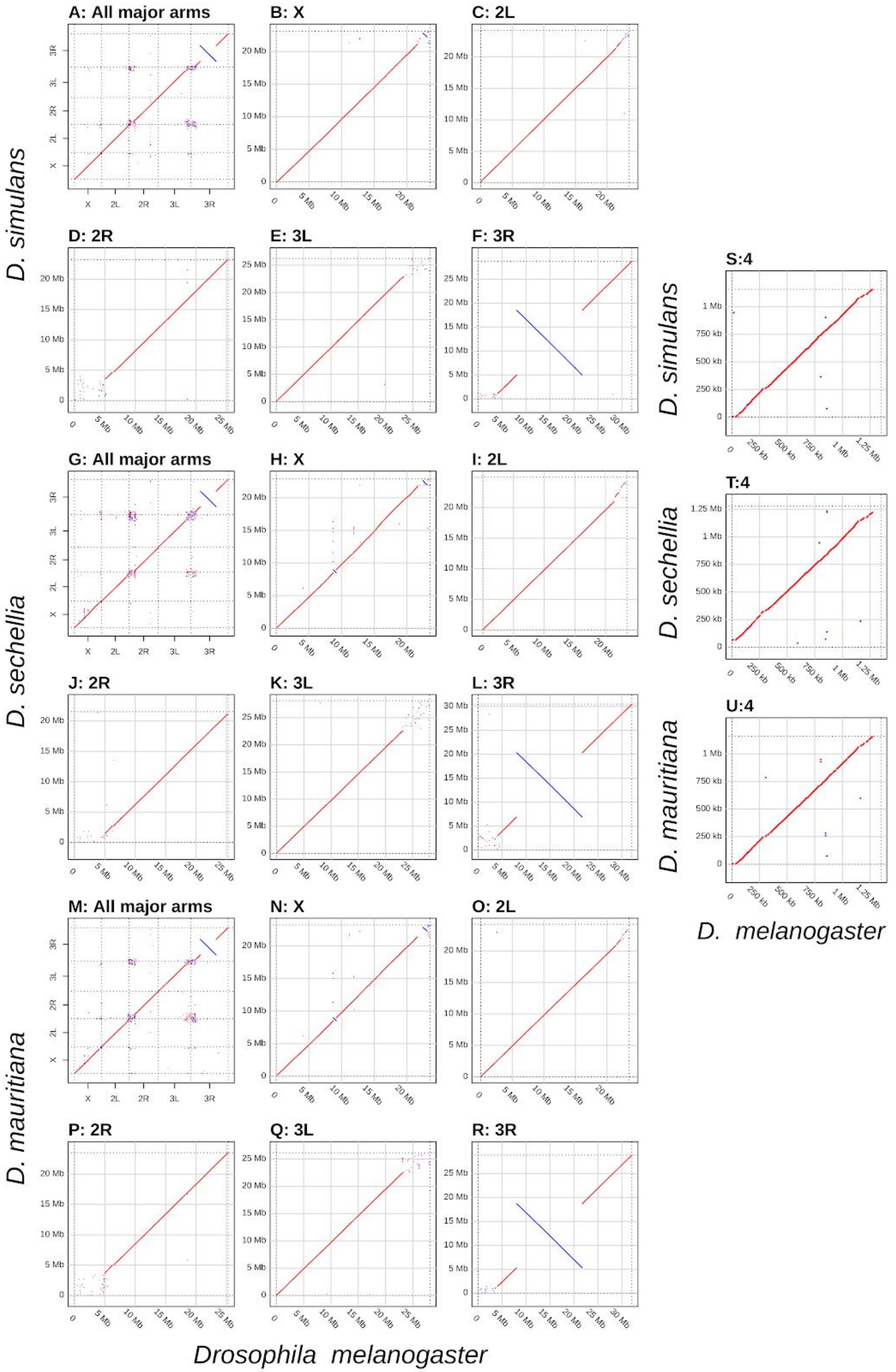
A: All major arms
B: X
C: 2L
D: 2R
E: 3L
F: 3R
G: All major arms
H: X
I: 2L
J: 2R
K: 3L
L: 3R
M: All major arms
N: X
O: 2L
P: 2R
Q: 3L
R: 3R
S:4
T:4
U:4
D. simulans
D. sechellia
D. mauritiana
Drosophila melanogaster
D. melanogaster

Figure S4. Dotplots for *D. melanogaster* against the members of the sim-complex species, following Figure 2B. In all plots, the X axis represents coordinates for *D. melanogaster* and the Y represents the sim-complex species. Panels A-F are *D. simulans*, G-L are *D. sechellia*, and M-R are *D. mauritiana*. The first panel for each species (A, G, M) represents the entire genome. The divisions in the dotplots span the five major Muller elements in order: A (X), B (2L), C (2R), D (3L), E (3R). The next 5 panels are zoomed into each Muller element: *D. simulans* spans panels B-F; *D. sechellia* spans panels H-L; *D. mauritiana* spans panels N-R. Dotplots of Muller F or Chromosome 4 are shown in S-U.

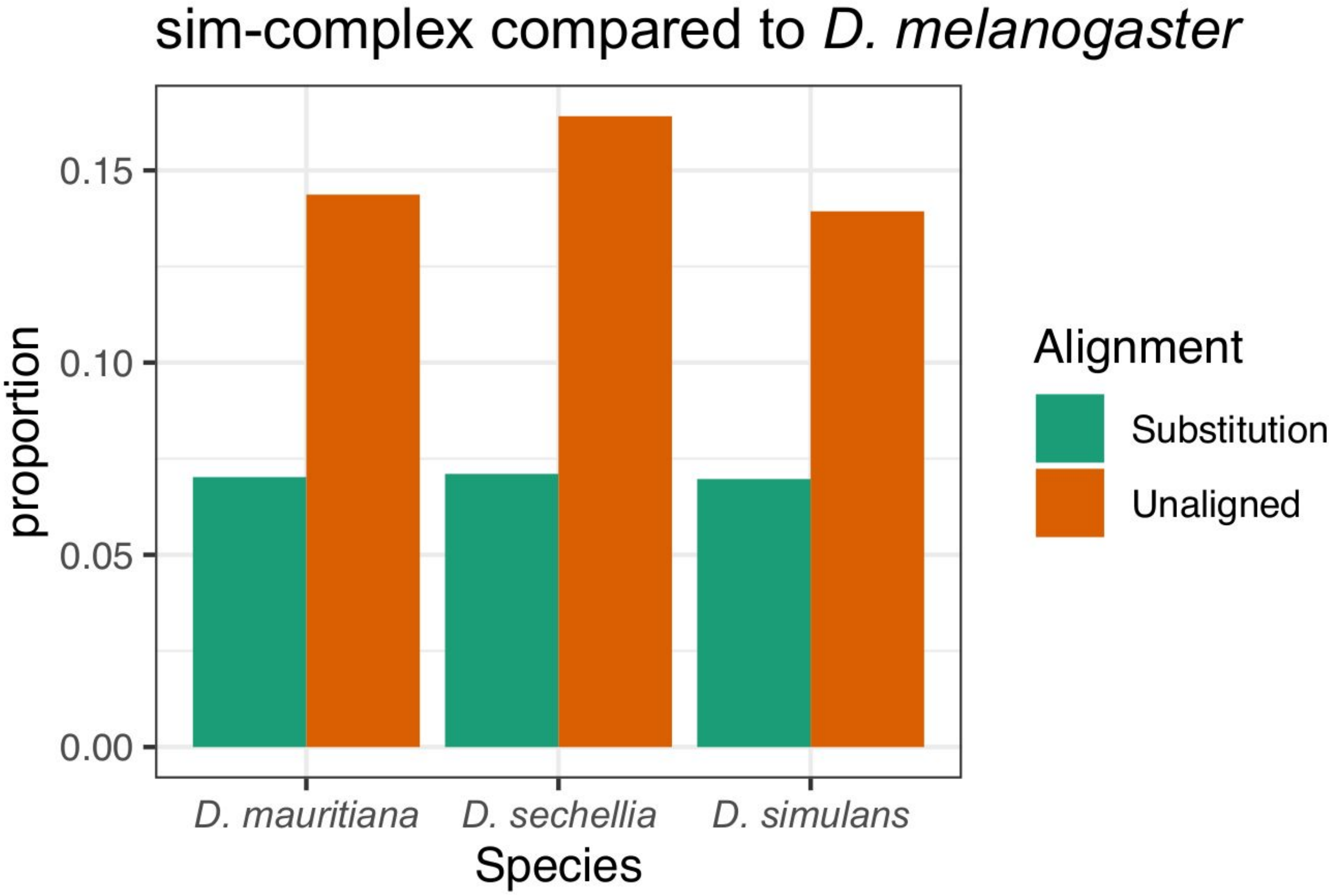


Figure S5. Proportion of scaffolded sim-complex species genomes that differ from *D. melanogaster* due to single nucleotide variants (SNVs) and small indels within 1:1 mapping syntenic regions (green) and due to failure to align (orange). Each sim-complex species genome was aligned to the reference *D. melanogaster* genome (Hoskins et al. 2015) using NUCmer (-g 200) and the sequence conservation and divergence was quantified with dnadiff (Marçais et al. 2018). Non-identical nucleotides between two genomes within unique alignments were counted as substitutions. The portion of the genomes that failed to align were counted as unaligned.

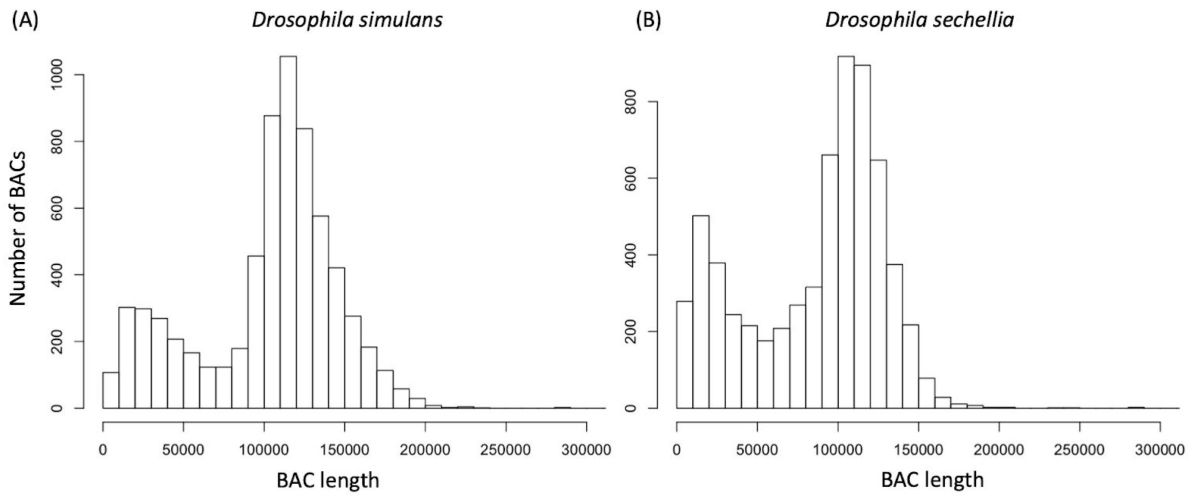


Figure S6. The size of BAC libraries in *D. simulans* (A) and *D. sechellia* (B). To validate the completeness and qualities of our assemblies. We mapped the two ends of each BAC to our assemblies to infer the length of BACs. The majority of BACs (>95%) are mapped into the same contigs with reasonable distance (< 300 kb).

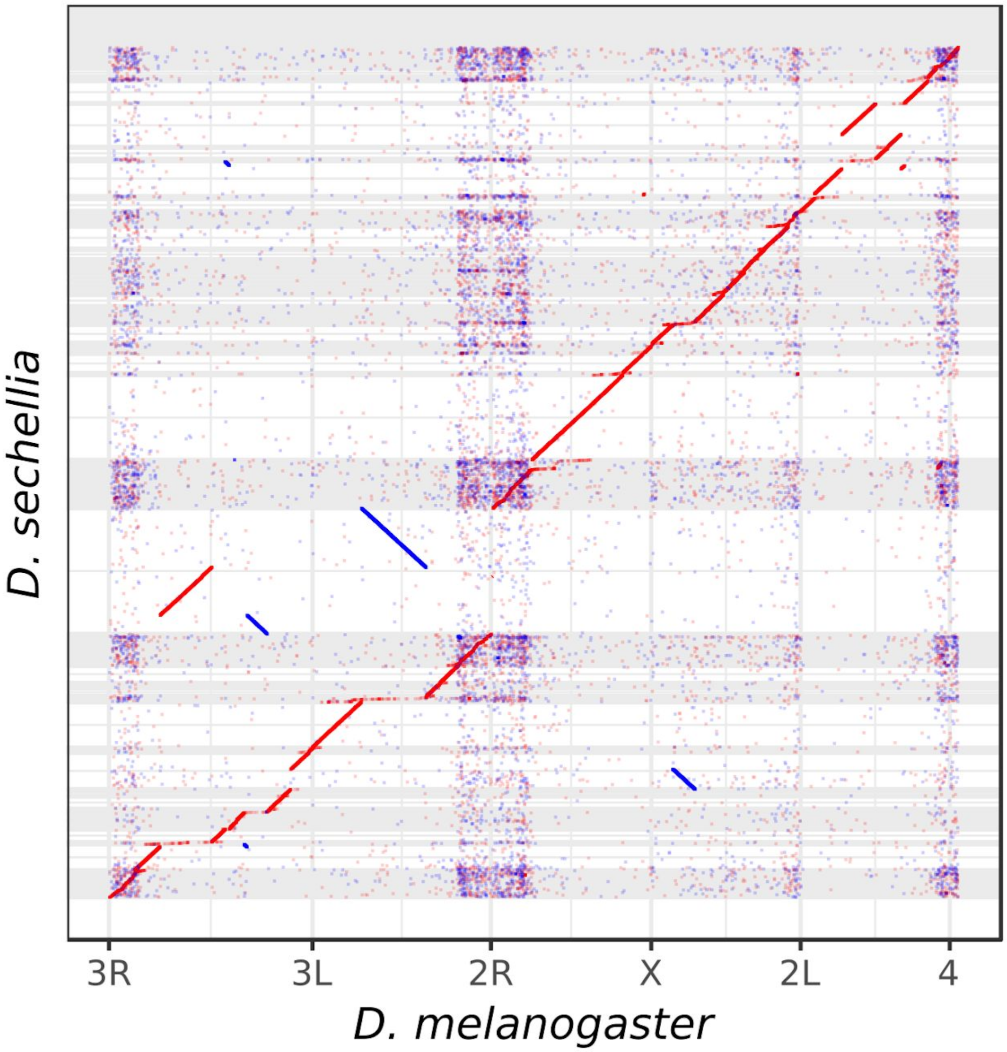


Figure S7. Dot plot between the FlyBase *D. sechellia* assembly (release 1.3) and *D. melanogaster* reference assembly. Except for the large inversion on 3R, all large scale off-diagonal and inverted alignments shown here are mis-assemblies in the previous *D. sechellia* assembly and are corrected in our *D. sechellia* assembly (Fig. 1, supplementary Fig. S4G-L) reported in this paper.

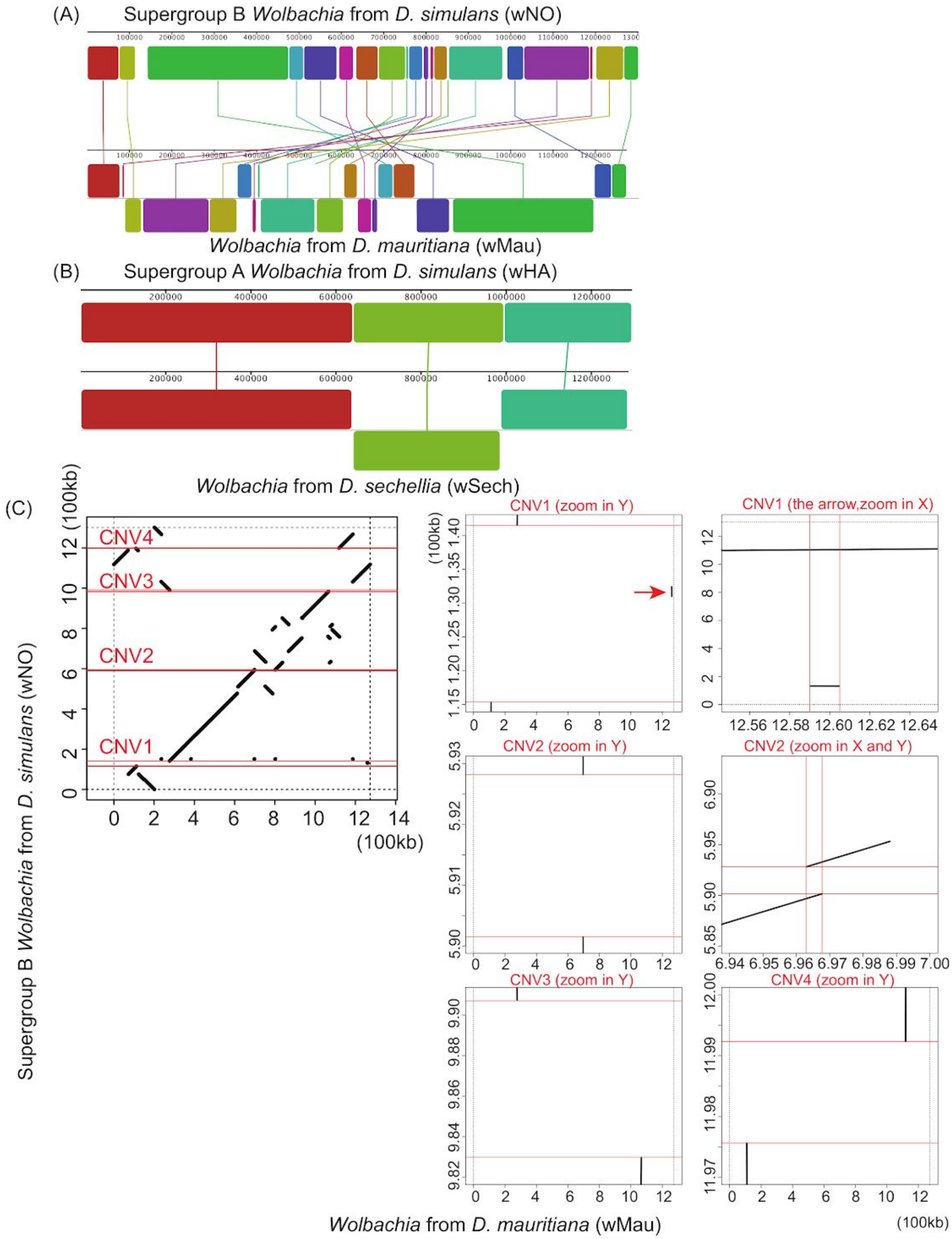


Figure S8. The genome alignment of *Wolbachia* sequences from *D. mauritiana* (*w*Mau) and *D. sechellia* (*w*Sech). We aligned *w*Mau to the supergroup B of *Wolbachia* reference from *D. simulans* (A), and aligned *w*Sech to the supergroup A of *Wolbachia* reference from *D. simulans* (B) using Mauve. Colored rectangles show positions of syntenic collinear blocks compared to the reference. The 4 CNVs between *w*No and *w*Mau previously identified by Meany et al. (Meany et al. 2019) are located on breakpoints of rearrangements (C). These deleted regions of the *w*Mau genomes include many candidate genes for loss of cytoplasmic incompatibility and minimal fecundity effects in *w*Mau, including many phage-related loci. The coordinate of *w*Mau and *w*Sech is adjusted based on the reference, *w*No and *w*Ha, respectively, and only represents the size of homology blocks.

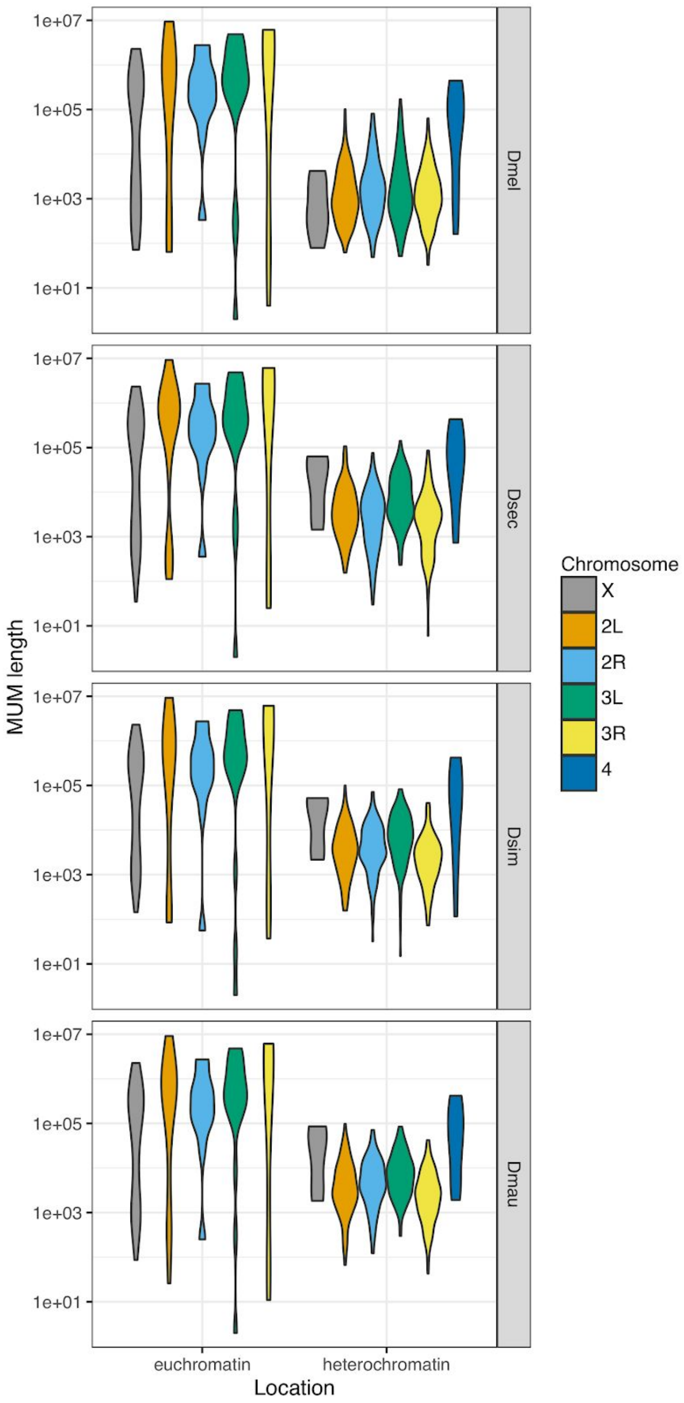


Figure S9. Length of collinear groups across mel-complex species. We extracted 540 one-to-one linear alignments (MUMs) in four species from Mauve's results and calculated their length. We then categorized alignments by their chromosome location. The length of MUMs are significantly shorter in heterochromatin than euchromatin for all autosomal arms (two-tail t test, P < 1e-8), but not for the X chromosome (two-tail t test, P > 0.18)

(A)

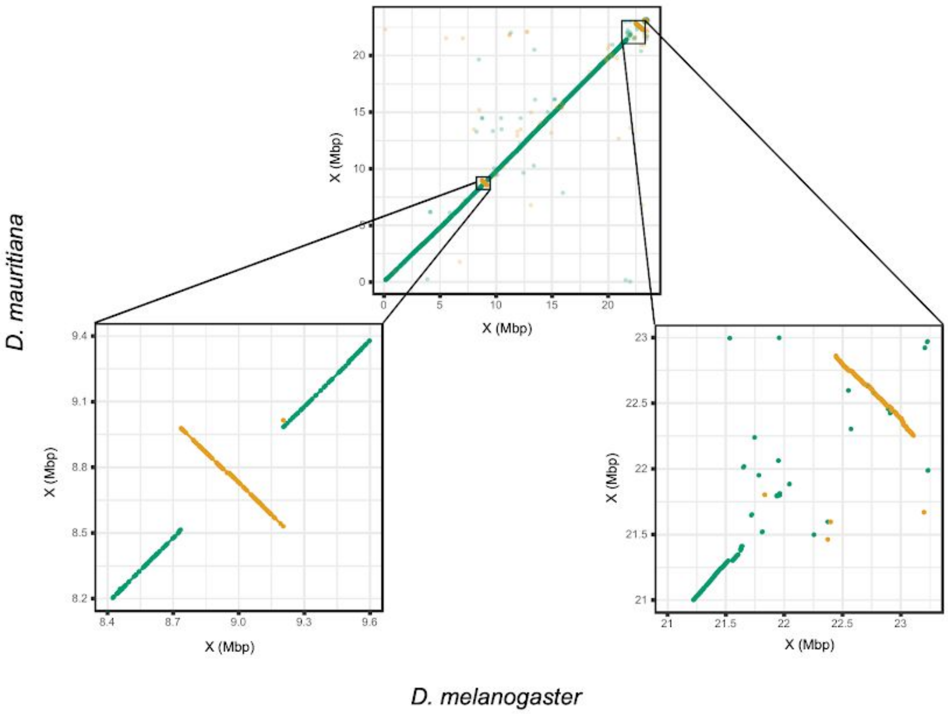

D. mauritiana
X (Mbp)
X (Mbp)
X (Mbp)
X (Mbp)
X (Mbp)
X (Mbp)
D. melanogaster


(B)

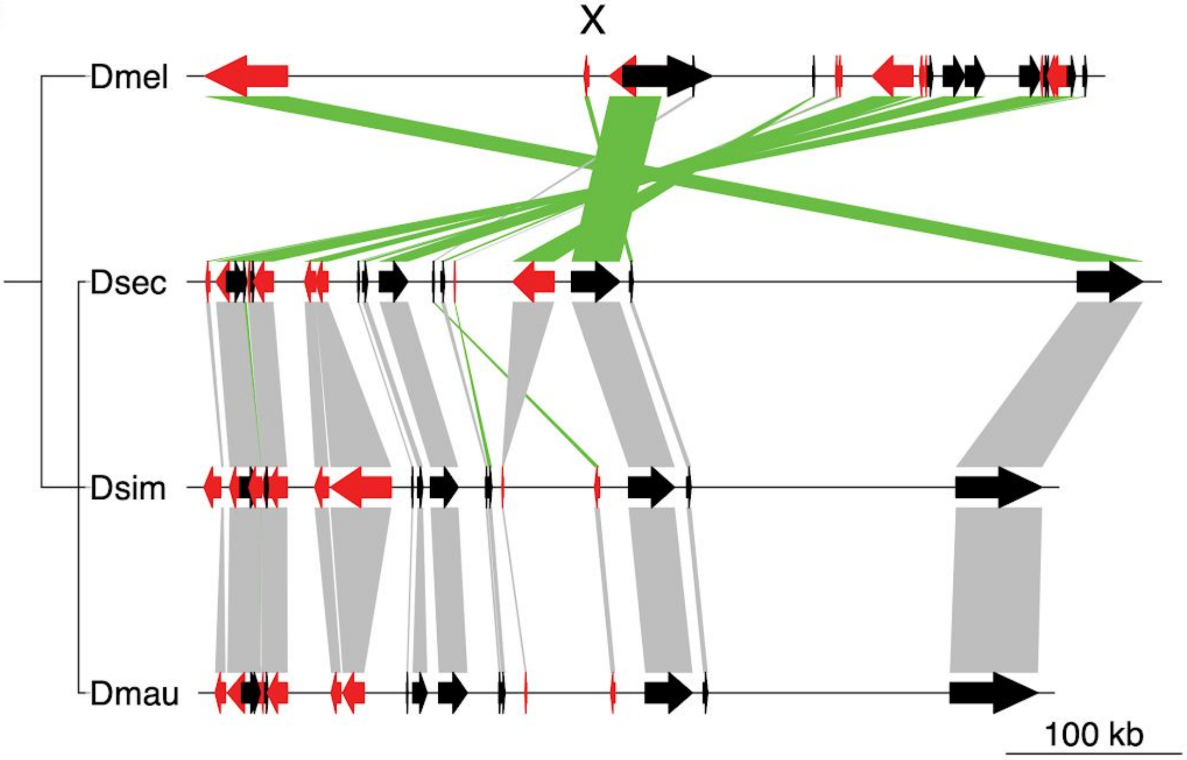

X
Dmel
Dsec
Dsim
Dmau
100 kb

Figure S10. Two large inversions (>100kb) on X chromosomes between *D. melanogaster* and the sim-complex clade species. (A) Dotplots between *D. mauritiana* and *D. melanogaster* showing two large inversions on the *D. mauritiana* X chromosome. The top figure is of the whole chromosome while the bottom two figures are magnified versions of regions in the top figure. The inversion in the middle of the chromosome is ~460kb and is also present in *D. sechellia* (not shown here). The forward alignments are in green and the inverted alignements are in yellow. (B) The inversion at the boundary of euchromatin and centromeric heterochromatin is ~700kb and is present in all three sim-complex species. The red and black arrows indicate orthologous genes among species with different orientation. The orothogs between species are connected by bold lines, and the lines are marked in green when the orientation changed between species

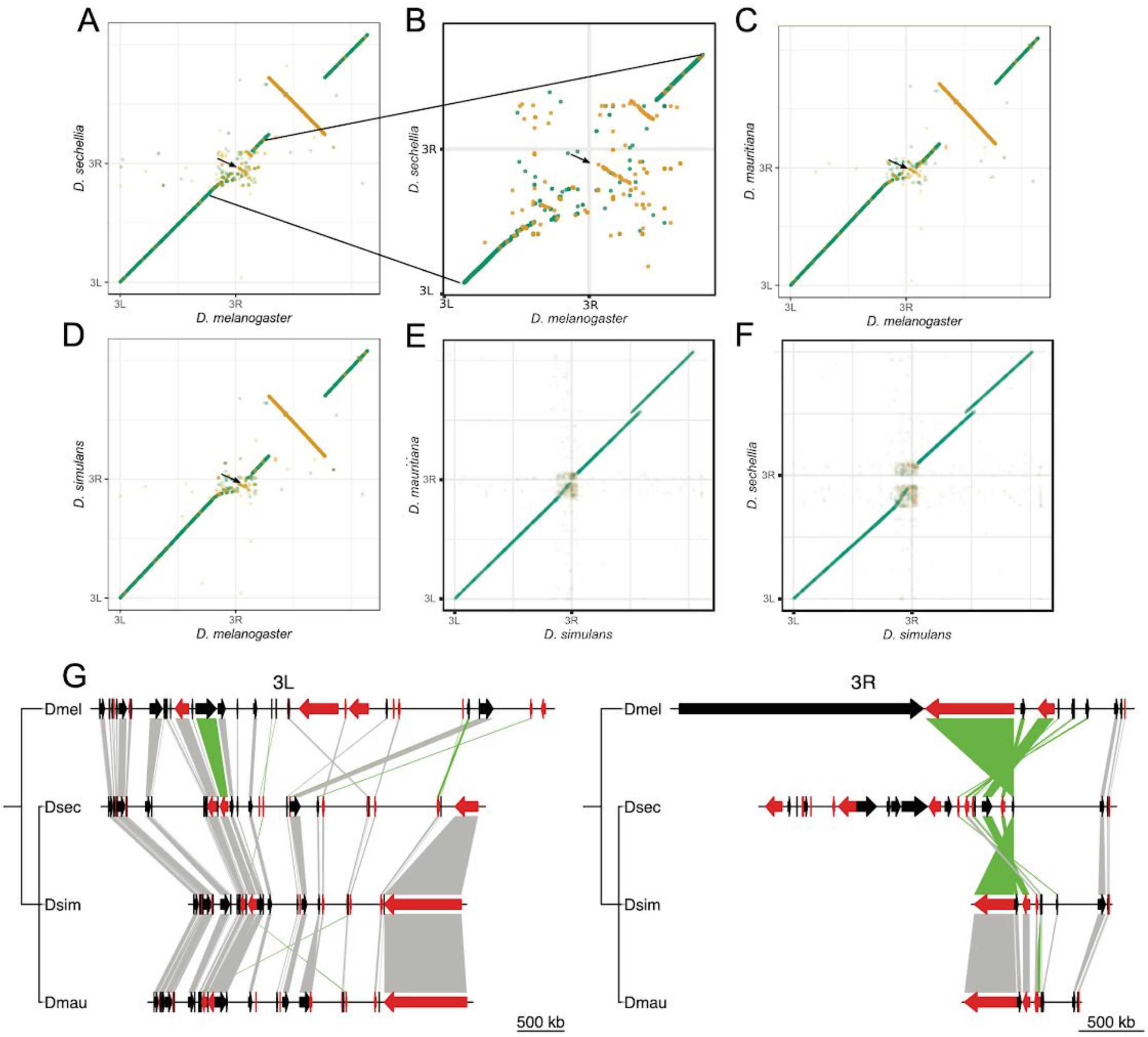

Figure S11. Inversions between sim-complex and *D. melanogaster* on Chromosome 3. (A–F) Alignment dotplots between sim-complex and *D. melanogaster* Chromosome 3. The MUMmer alignment between the species genomes were filtered using delta-filter -l 200 and -l 2000 for the alignments involving and excluding *D. melanogaster*, respectively, to remove alignments due to repeats. The arrows point to a distal 3L pericentromeric segment in the sim-complex species (Y-axis) that maps to distal pericentromeric region of 3R in *D. melanogaster*. The breakpoint of the inversion proximal to centromere is not present in the assembly and therefore unknown. A. Alignment between *D. sechellia* and *D. melanogaster* Chromosome 3. B. The alignments in the pericentromeric region from A is expanded to show the precise location of the inversion. C. Alignment between *D. mauritiana* and *D. melanogaster* Chromosome 3. D. Dot Plot between *D. simulans* and *D. melanogaster*. E. Dot plot between *D. simulans* and *D. mauritiana*, showing the absence of the inversion. F. Dot plot between *D. simulans* and *D. sechellia*, which is also missing the inversion. The green alignments represent sequences on the Y-axis that are in the same orientation as the sequences in the X-axis, whereas the yellow alignments are in the inverted orientation with respect to the X-axis genome. (G) The gene synteny of heterochromatic genes on Chromosome 3 among four species. The red and black arrows indicate orthologous genes among species with different orientation.The orthologs between species are connected by bold lines, and lines are marked in green when orientation differs between species.

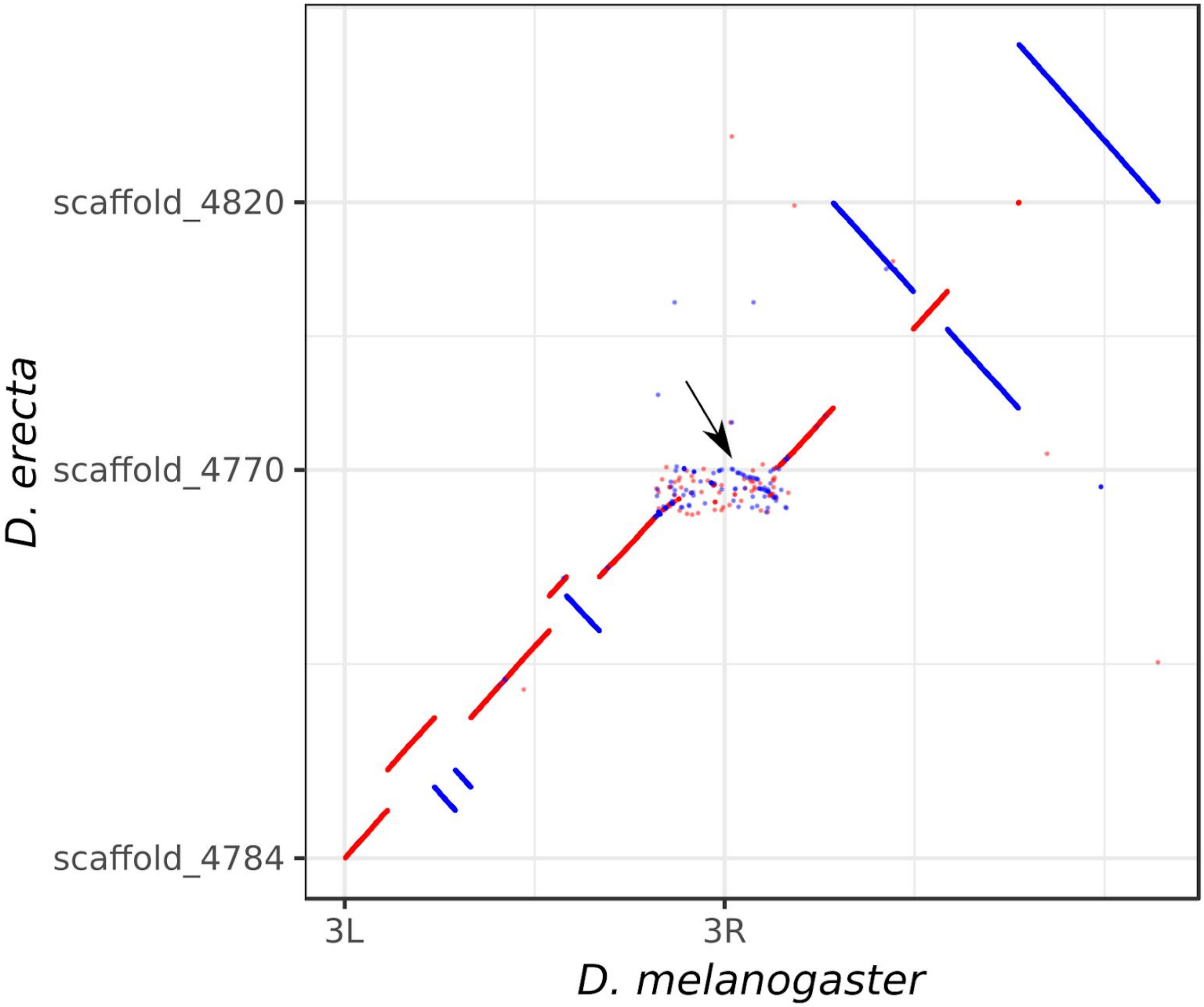


Figure S12. Dot plot of the alignment between the *D. melanogaster* and the the *D. erecta* Chromosome 3 scaffolds. As shown by the arrow, a part of the *D. erecta* 3L pericentric heterochromatin maps to the 3R pericentric heterochromatin of *D. melanogaster*. This inversion is the same as the inversion shown in Fig. S9, suggesting that the inversion occurred in the melanogaster lineage.

(A)

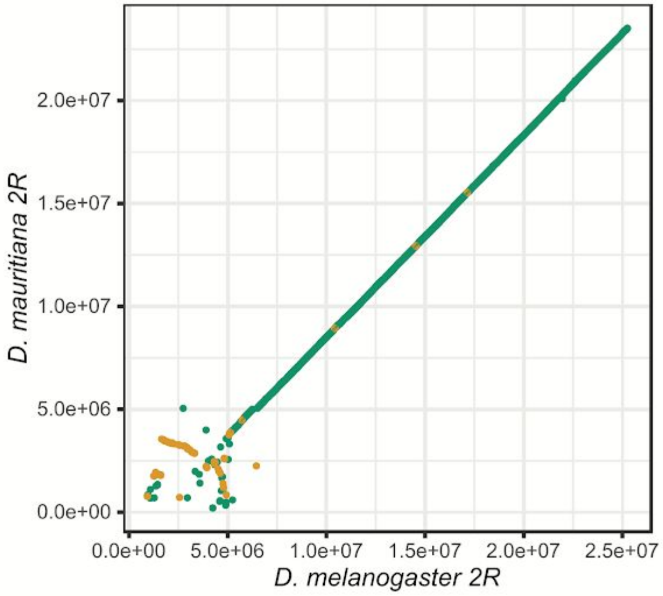


(B)

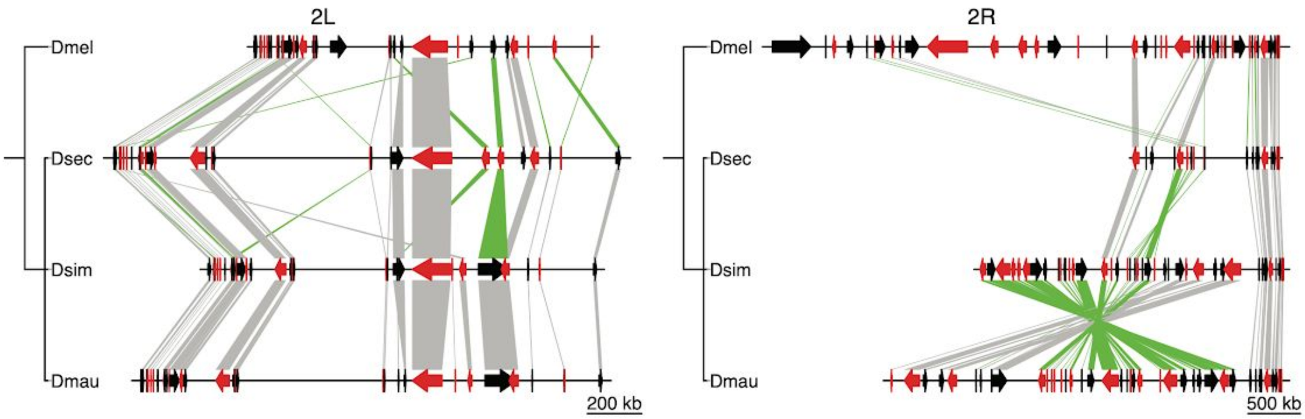


Figure S13. A large species-specific heterochromatic inversion on *D. mauritiana* 2R (A) Dot plot between *D. melanogaster* and *D. mauritiana* 2R chromosome arm showing the presence of an inversion inside the pericentromeric heterochromatin. The alignments that are in the same orientation as the *D. melanogaster* sequence are in green and the inverted alignements are in yellow. (B) The gene synteny of heterochromatic genes on the second chromosome among four species. The red and black arrows indicate orthologous genes among species with different orientation. The orthologs between species are connected by bold lines, and the lines are marked in green when the orientation changes between species.

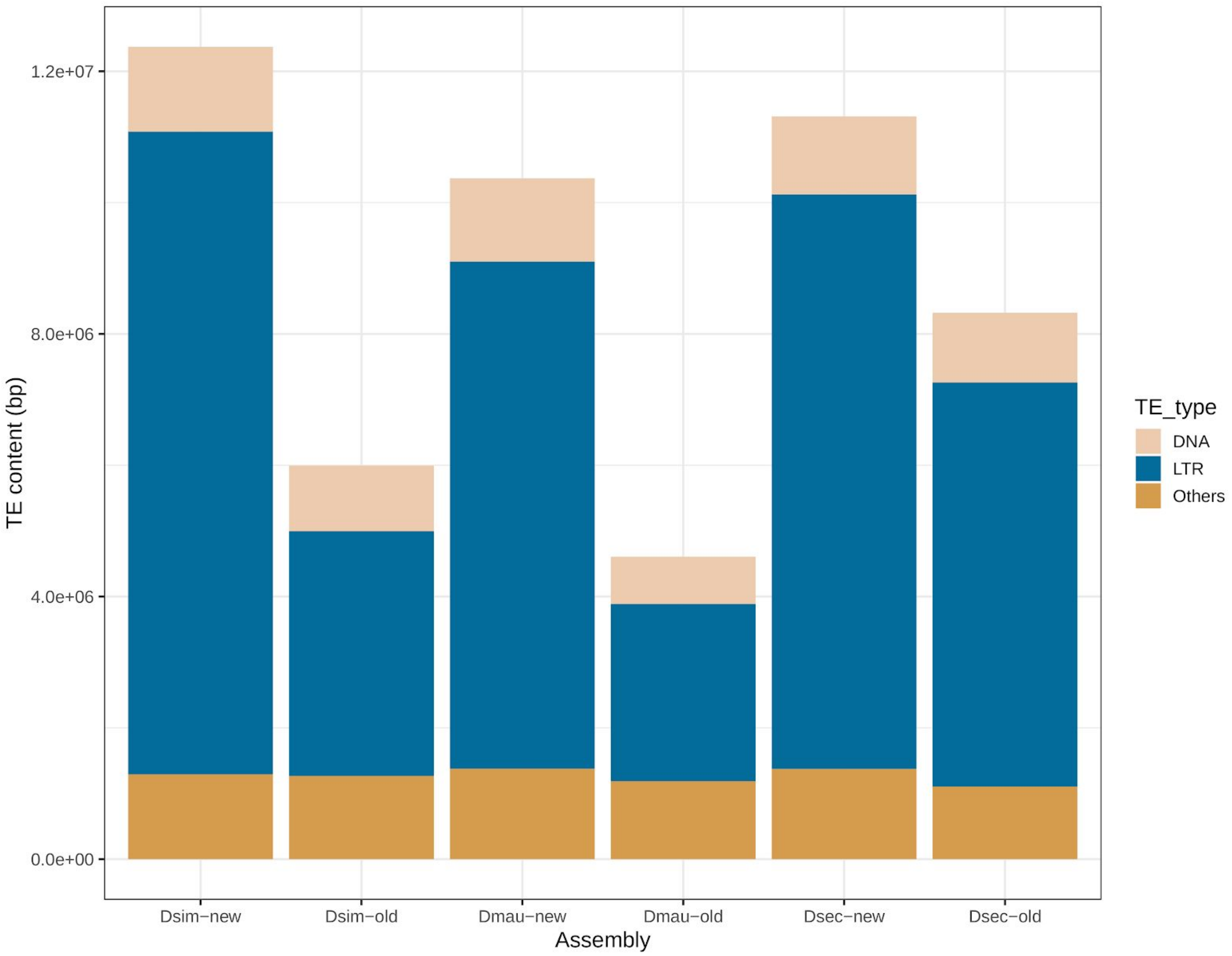


Figure S14. Total amount of LTR retrotransposons, DNA transposons and others (including non-LTR retrotransposons) in the sim-complex assemblies reported here and the sim-complex assemblies published previously (Hu et al. 2013; Garrigan et al. 2012; *Drosophila* 12 Genomes Consortium 2007).

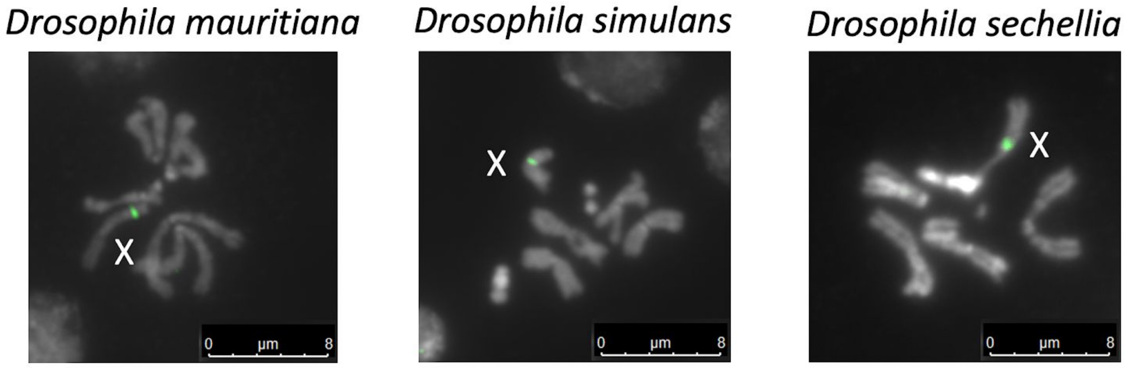


Figure S15. Fluorescence *in situ* hybridization of new X-linked satellite (193XP: Green) on mitotic chromosomes. We dissected the brains from third instar larva and used probes to detect repetitive sequences by fluorescence in situ hybridization in the sim-complex species.

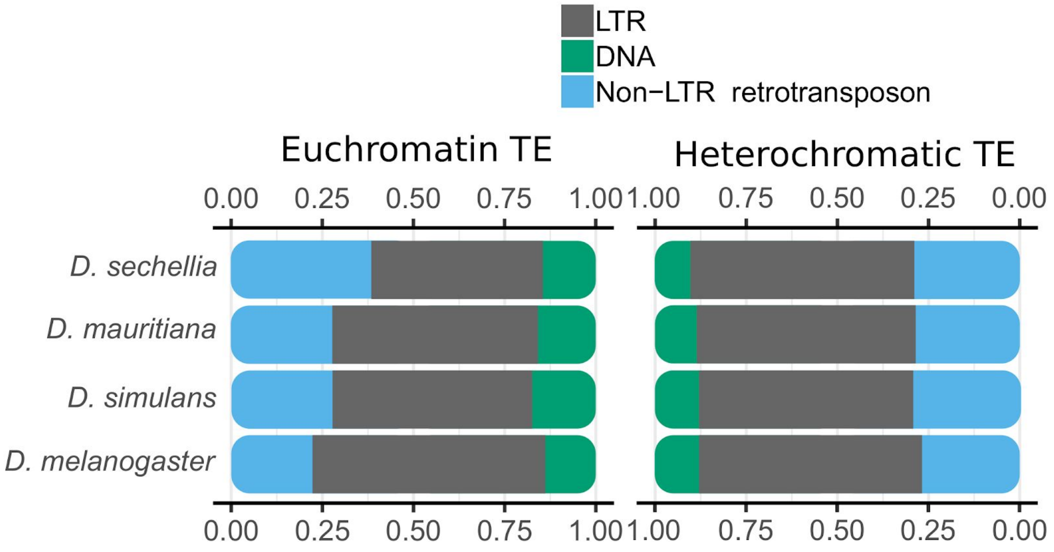


Figure S16. The proportion of different transposons in euchromatic and centromeric heterochromatin regions in the mel-complex species. The proportions appear more similar across species in the heterochromatic regions than they are in the euchromatic regions.

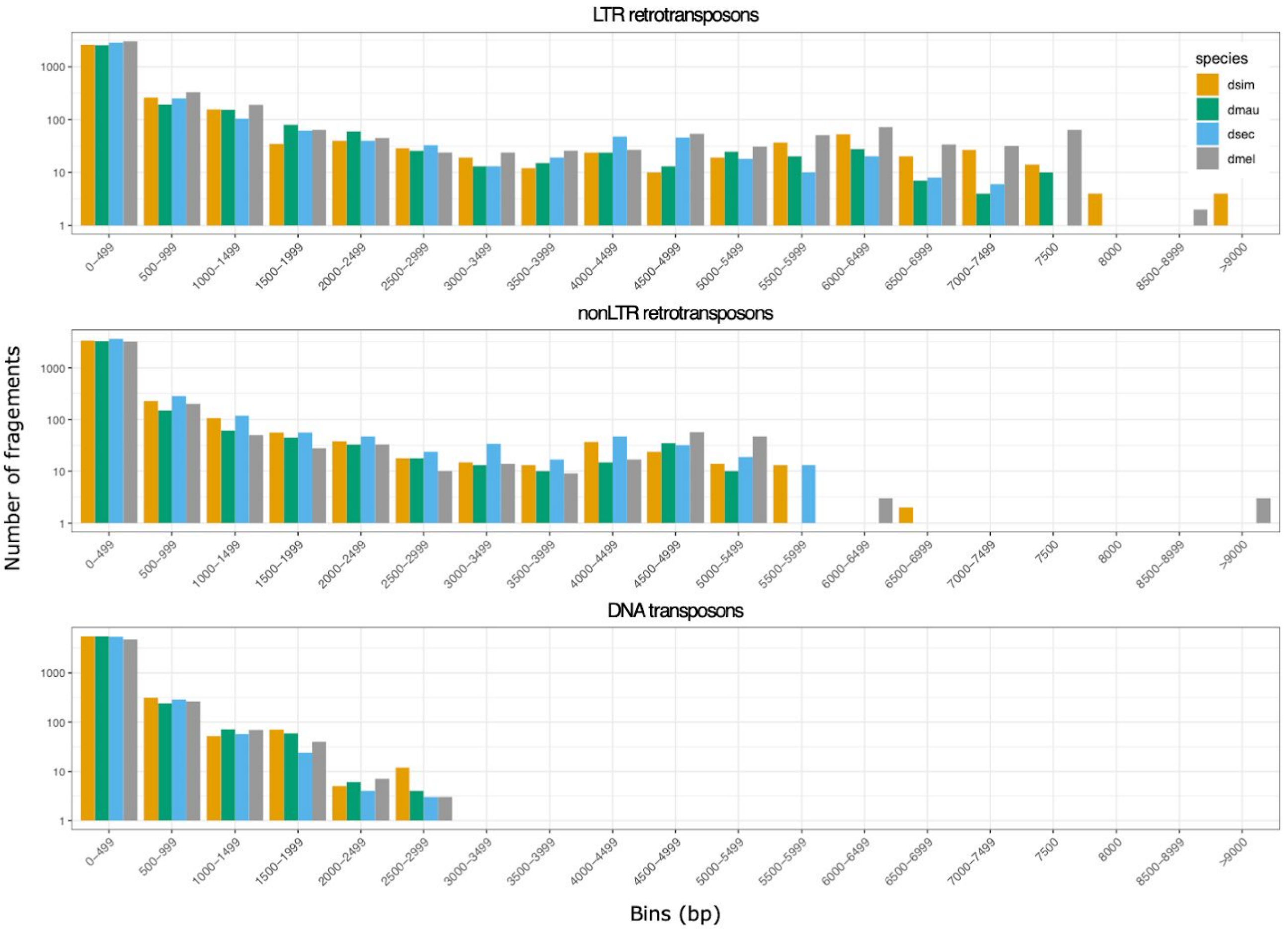


Figure S17. The distributions of transposon sequence lengths in the four mel-complex species. We classified the TEs by LTR and Non-LTR retrotransposons and DNA transposons.

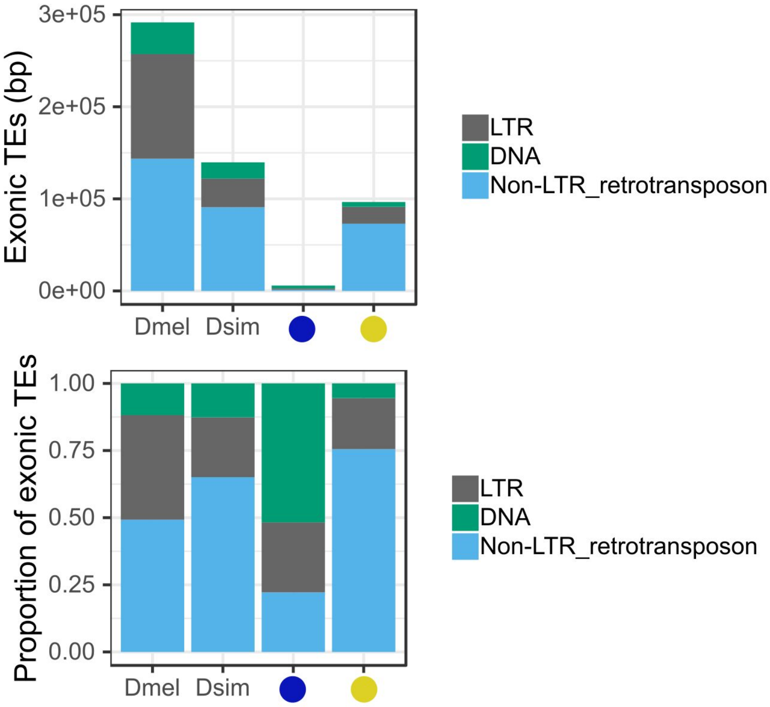


Figure S18. TE content in the exons of 6,984 genes annotated by Iso-Seq in *D. simulans,* their homologs in *D. melanogaster*, and the orthologous segments in the ancestral branches of the sim-complex and mel-complex. Bar labels follow Figure 4: the ancestral lineages of the sim-complex species are represented by blue dots and the mel-complex species are represented by yellow dots. The top panel shows the total number of bases and the bottom panel shows the relative proportion.

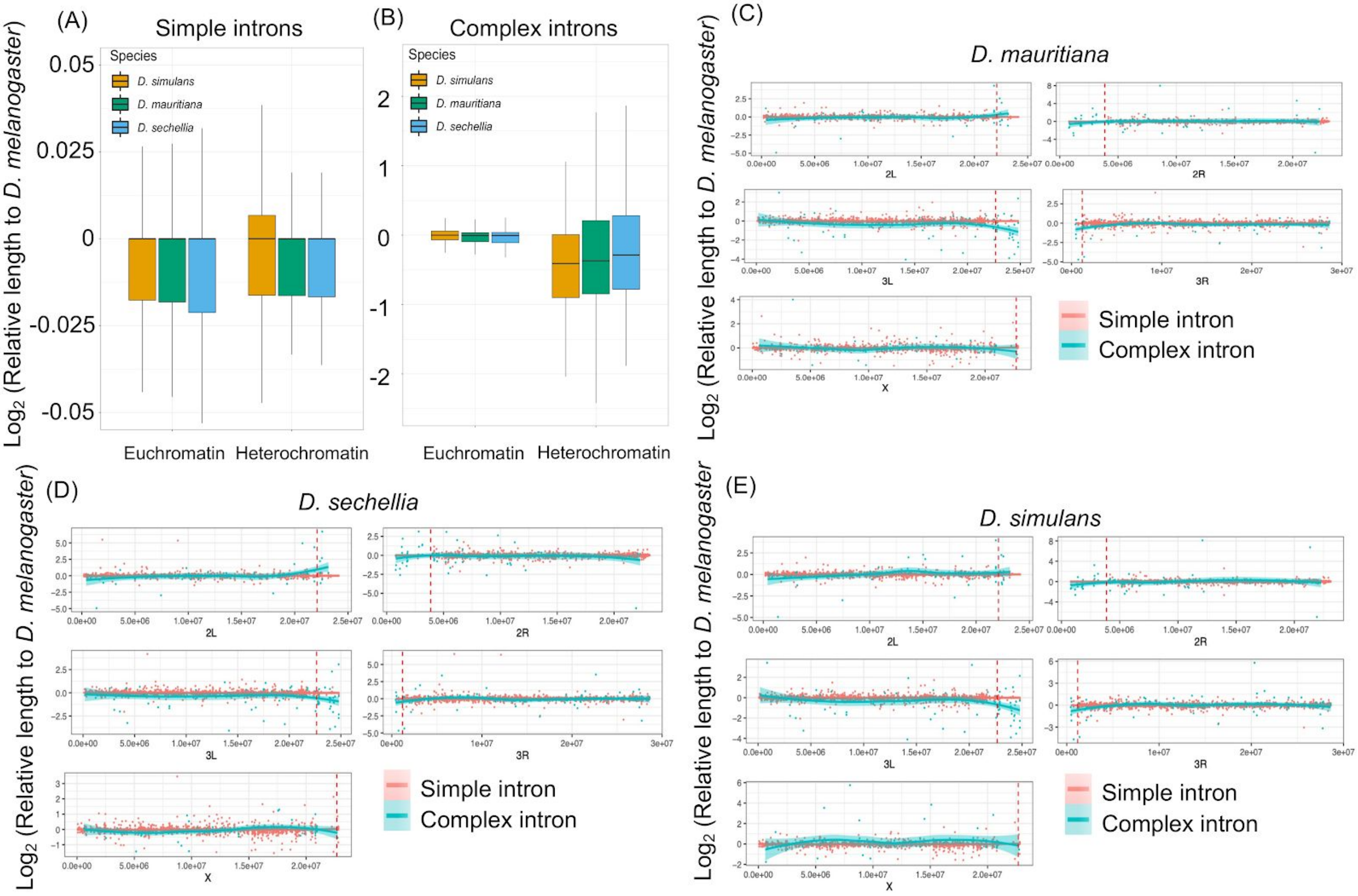


Figure S19. The intron length changes in the sim-complex species relative to *D. melanogaster*. We classified the introns based on whether they contained repetitive sequences (transposons or complex satellites). The introns without repetitive sequences (simple introns) are shown in (A) and the introns with repetitive sequences (complex introns) are shown in (B). (C-E) Changes in the intron lengths across the genome based on *D. simulans* coordinates. Each dot represents an intron and the lines are the LOESS regression curves for the changes and chromosome coordinates. Red points and lines indicate the simple introns and blue points and lines indicate the complex introns. The red dot lines represent the boundaries between euchromatin and heterochromatin.

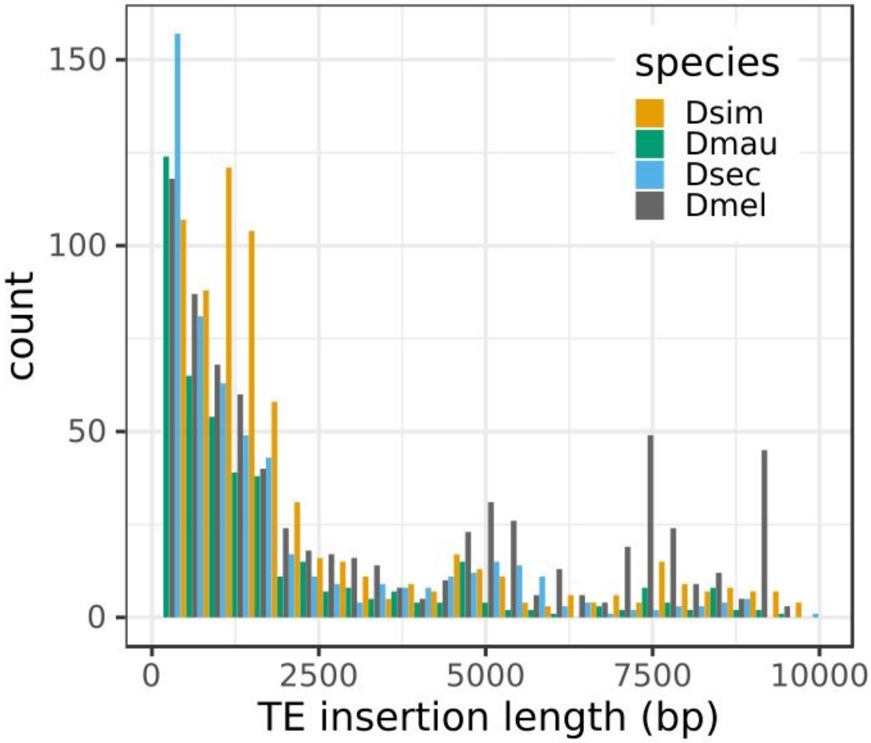


Figure S20. The distribution of intronic transposon insertion lengths in the mel-complex species.

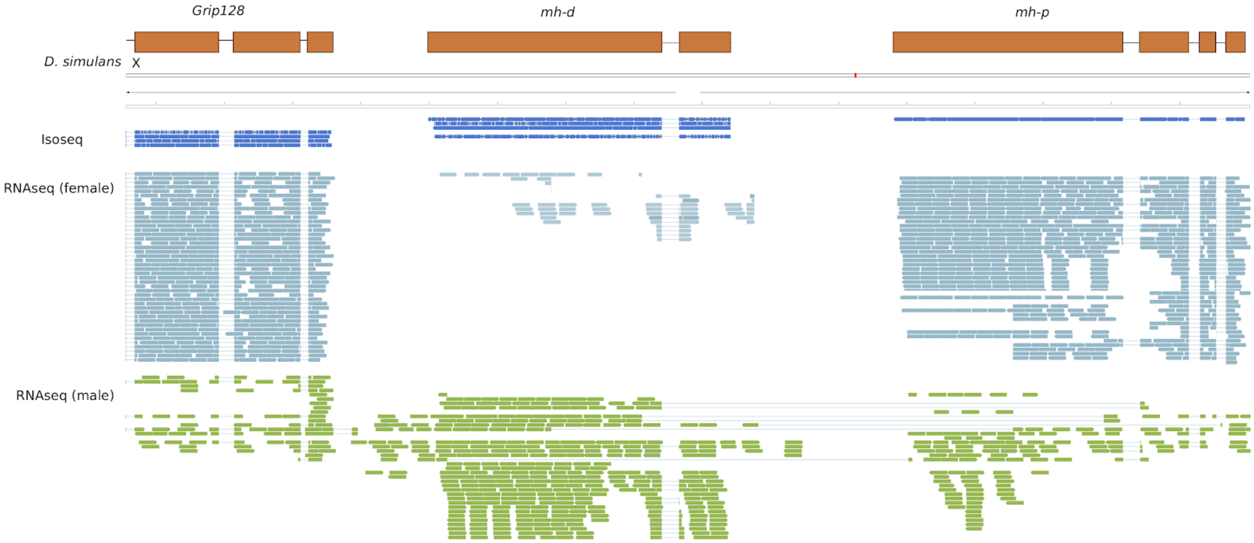


Figure S21. Iso-Seq and Illumina transcriptome reads mapped to copies of *maternal haploid* (*mh*) in *D. simulans*. The *mh-d* copy shows male-biased expression whereas *mh-p* copy shows female-biased expression.

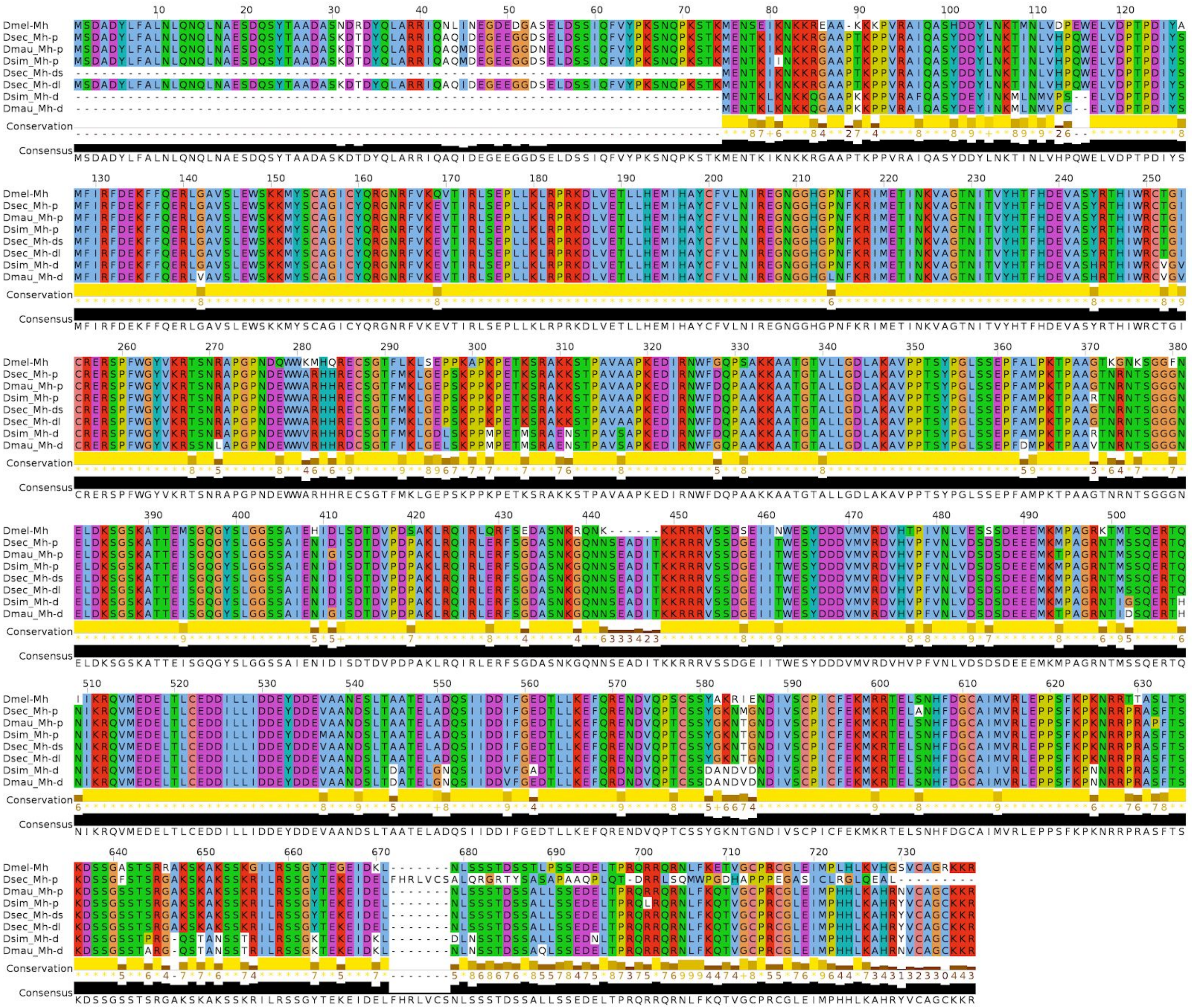

Figure S22. Alignment of proteins encoded by *D. melanogaster mh* and sim-complex *mh-p* and *mh-d* copies. As expected from the gene annotation, MH-p proteins are bigger than the MH-d proteins. *D. sechellia* has three copies of the *mh* gene but the copy showing similar size to the MH-d in the other two sim-complex species do not share all derived substitutions present in the MH-d protein in *D. mauritiana* and *D. simulans* . The colors and numbers in the conservation track are a conservation index calculated by jalview (Waterhouse et al. 2009): paler colors and higher numbers indicate greater conservation of the physicochemical properties of the residues at those positions.

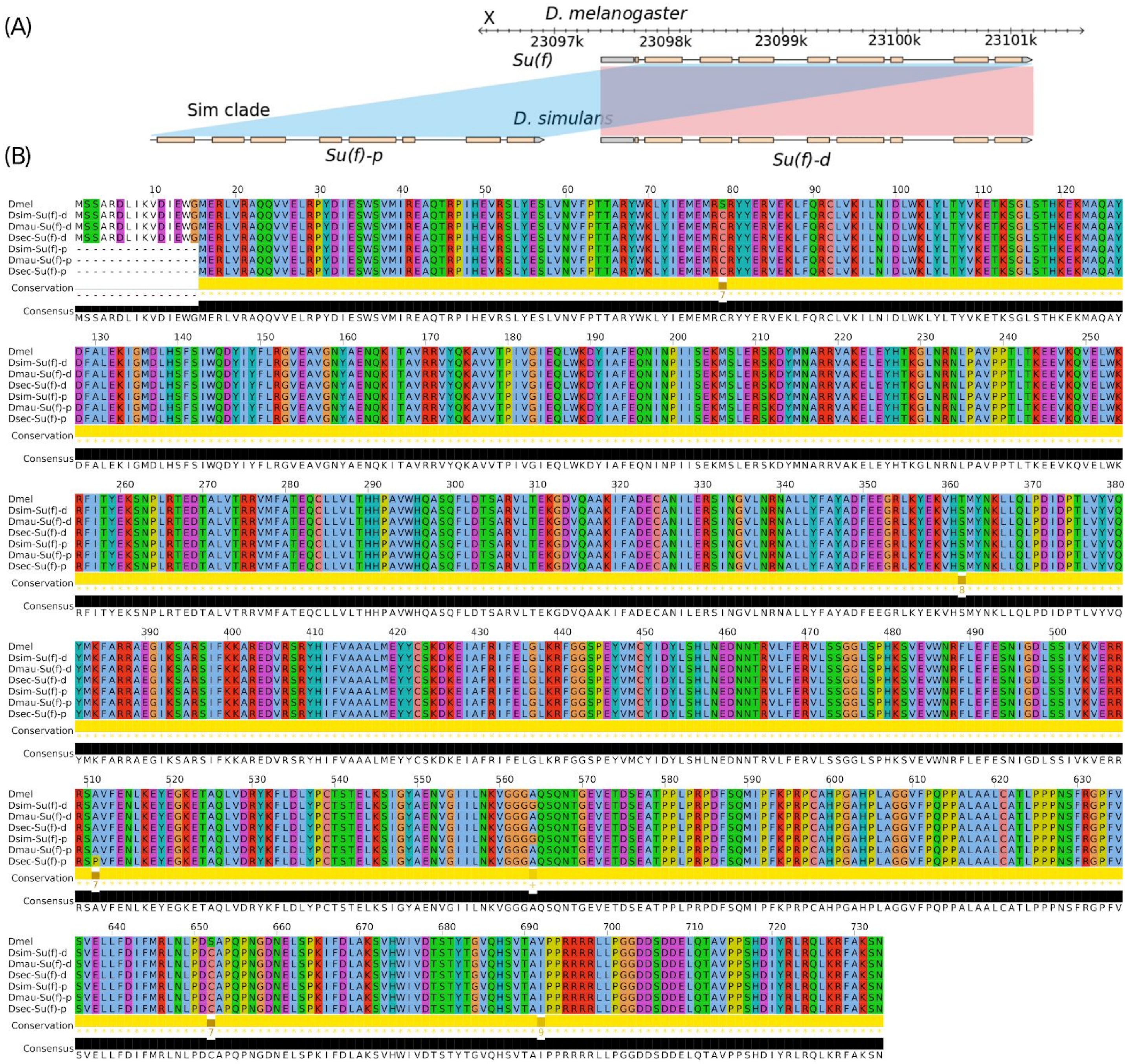


Figure S23. A. The duplication of *suppressor of forked* (*su(f)*) gene in the sim-complex. B. Protein sequence alignment of the *su(f)* copies showing the high sequence identity among the sim-complex species. The longest translated protein from the partial su(f) copy (*Su(f)-p*) was used for the alignment. The color codes and numbers in the conservation track are same as in Fig. S22.

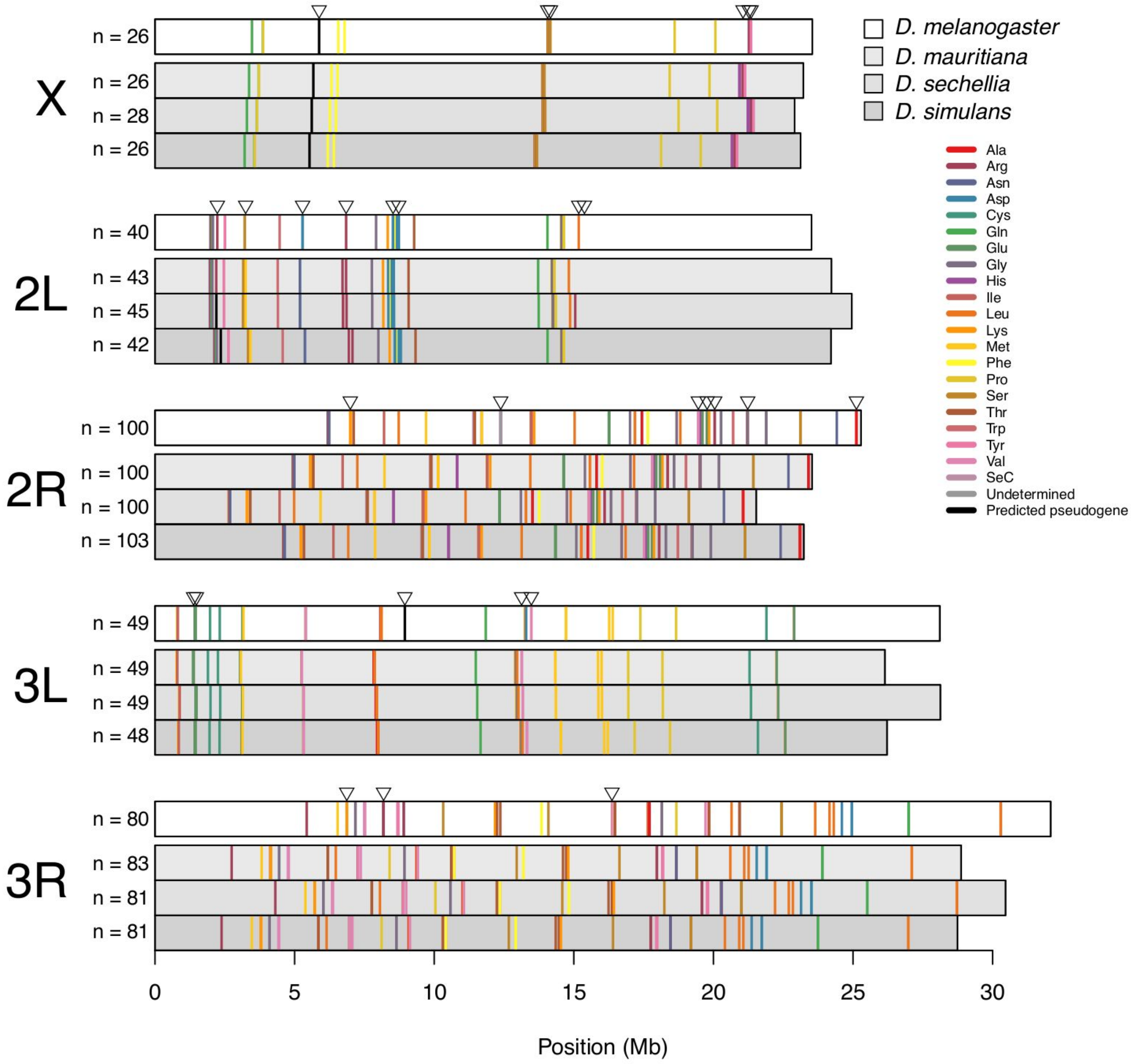


Figure S24. Distribution of all nuclear tRNAs annotated using tRNAscan-SE (Lowe and Eddy 1997) along Chromosomes X, 2L, 2R, 3L, and 3R in *D. melanogaster*, *D. mauritiana*, *D. sechellia*, and *D. simulans*. Chromosome lengths vary according to the assembly of each species and colored bars show the positions of different tRNA isotypes. Inverted triangles above the *D. melanogaster* track (top) correspond to the blocks in Figure 6A.

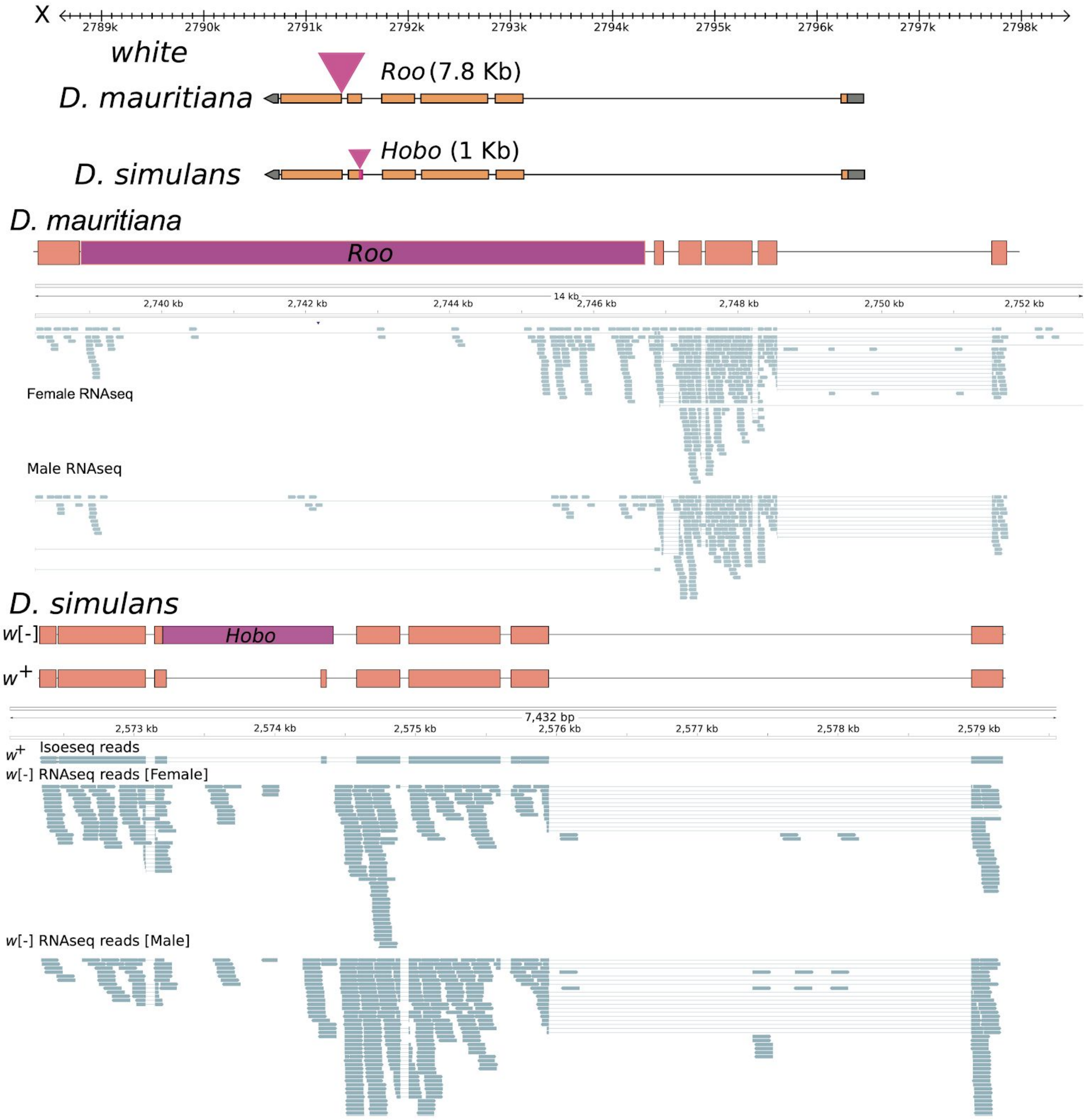


Figure S25. The transposon insertions disrupting the *white* gene of the two white-eyed sequenced strains: *D. mauritiana* (*w*12) and *D. simulans* ($w^{XD1}$).

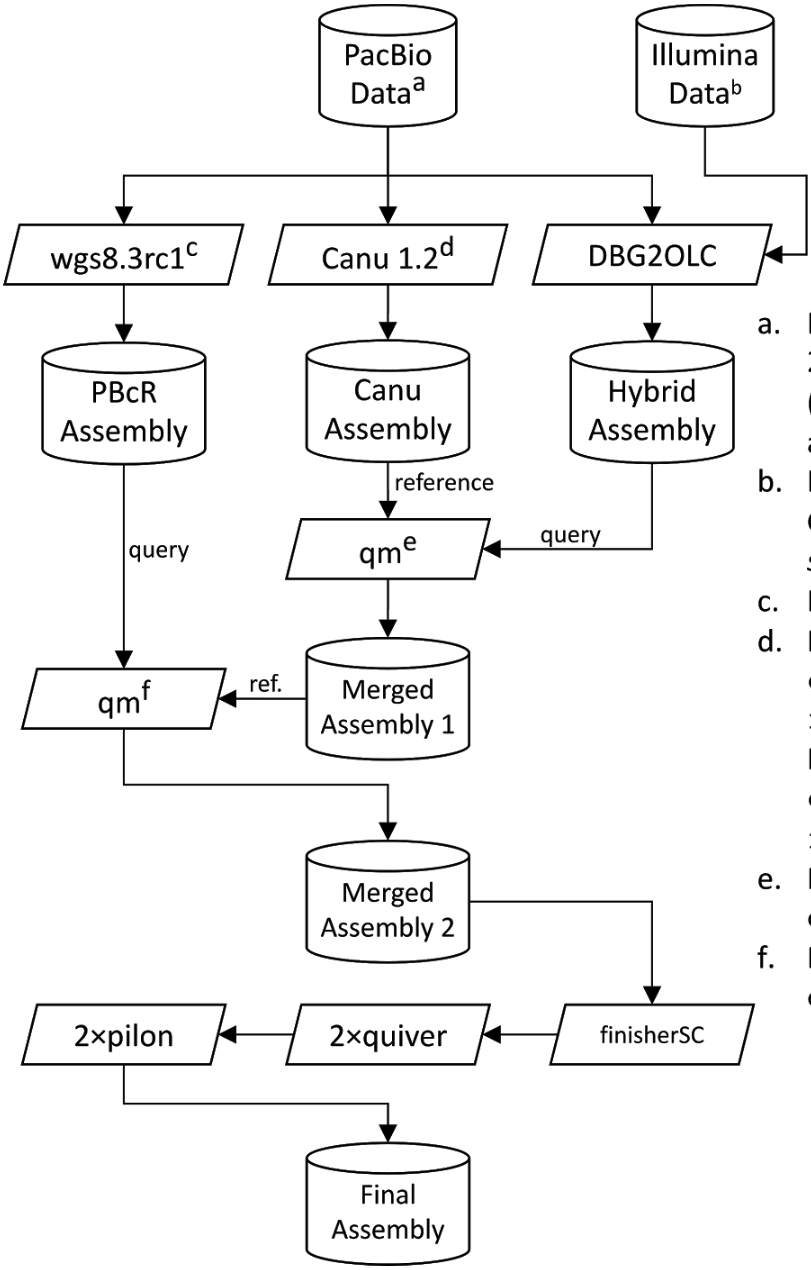


a. PacBio long reads: 21.5 Gb (NR50 = 14.9kb), 20.9 Gb for (NR50 = 15.7 kb), and 15 Gb for (NR50 = 15.2 kb) for *D. mauritiana*, *D. simulans*, and *D. sechellia*, respectively.
b. Illumina reads: 19.4 Gb, 17.68 Gb, and 20.65 Gb for *D. mauritiana*, *D. simulans*, and *D. sechellia*, respectively
c. Parameter: `-sensitive`
d. Parameters for *D. mauritiana* and *D. sechellia* : `genomeSize = 160m useGrid = false errorRate = 0.025`
   Parameters for *D. simulans:* `genomeSize = 160m useGrid = false errorRate = 0.035`
e. Parameters for quickmerge: `c=1.5, hc0=5.0, l=1000000`
f. Parameters for quickmerge: `c=1.5, hco=5.0, l=5000000`

Figure S26. The overview of our genome assembly approach. For each species, we generated three assemblies: a hybrid assembly with DBG2OLC and long read only assemblies with PBcR and Canu. The DBG2OLC assembly was constructed with the longest 3.9 Gb from each long read dataset and Illumina reads for each species. We ran PBcR (Berlin et al. 2015) using wgs8.3rc1 with the –sensitive parameter. We ran Canu 1.2 with genomeSize = 160m useGrid = false errorRate = 0.025 parameters for *D. mauritiana* and *D. sechellia* and with genomeSize = 160m useGrid = false errorRate = 0.035 parameters for *D. simulans* (https://github.com/marbl/canu/tree/5bd4744ad89b71243c7e52446c156956bd75672e; (Koren et al. 2017)). First, we merged the Canu and hybrid assemblies for each genome using quickmerge (Chakraborty et al. 2016; Solares et al. 2018) with the Canu assembly as the reference. We then used the merged assembly from the first step and merged it with the PBcR-sensitive assembly, with the former serving as the reference. We further improved the assemblies with finisherSC (Lam et al. 2015) with the raw reads. We polished all assemblies twice with Quiver (Chin et al. 2013), followed by final polishing with pilon (Walker et al. 2014). We then manually curated 10 misassemblies (supplemental Table S16; supplemental Fig. S27), including fixing the mitochondrial and Wolbachia genomes.

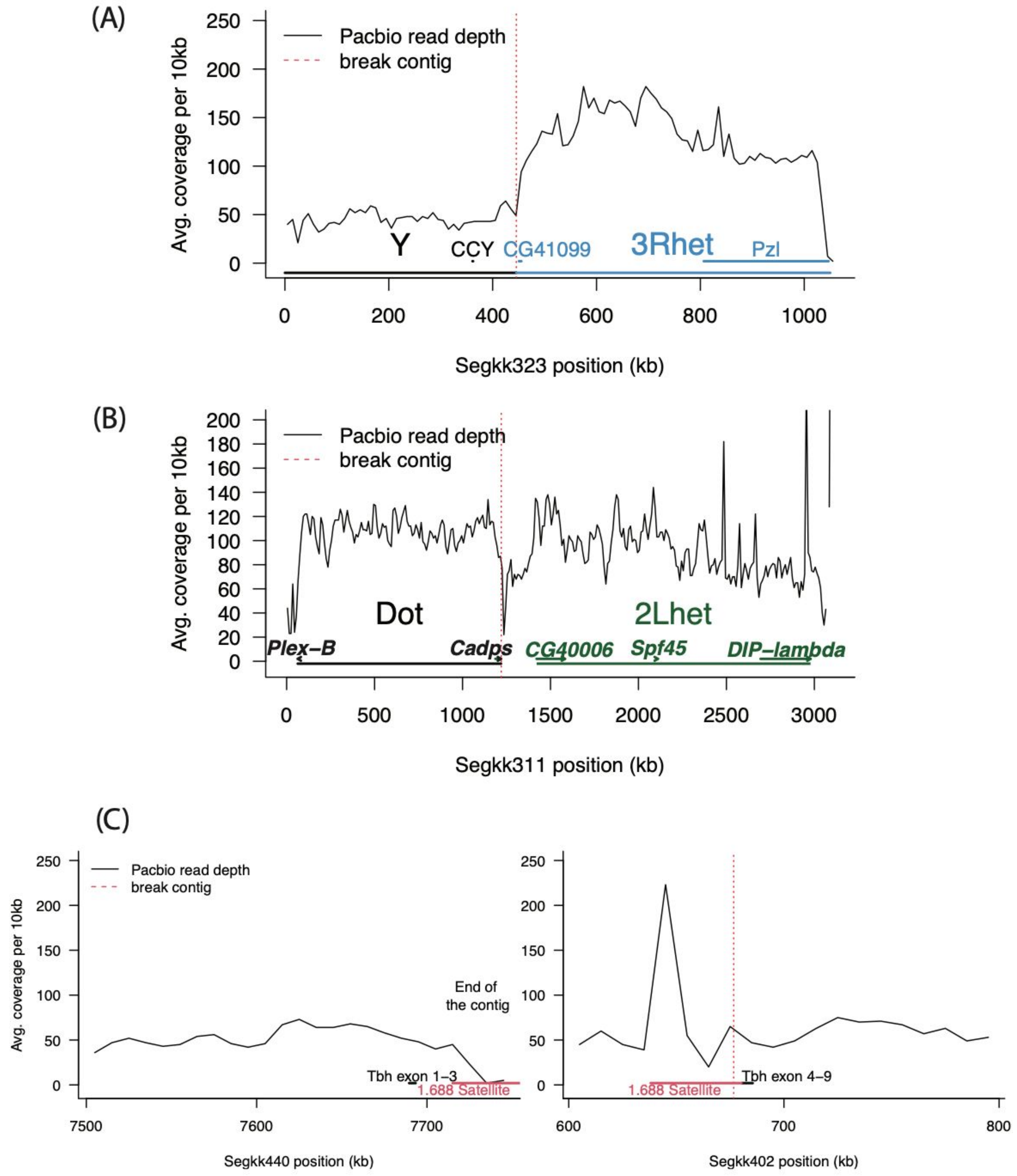


Figure S27. Correction of the misassemblies. We searched for the inconsistency of gene structure and PacBio read coverage across the genome and identified regions with misassemblies that fused two regions from different chromosomes (A and B) or caused disruption of a gene (C). We then broke the contigs in possible misassembled regions. We also found some regions lost or duplicated (e.g. mitochondrial sequences) during the assembling or reconciling assemblies (not shown).

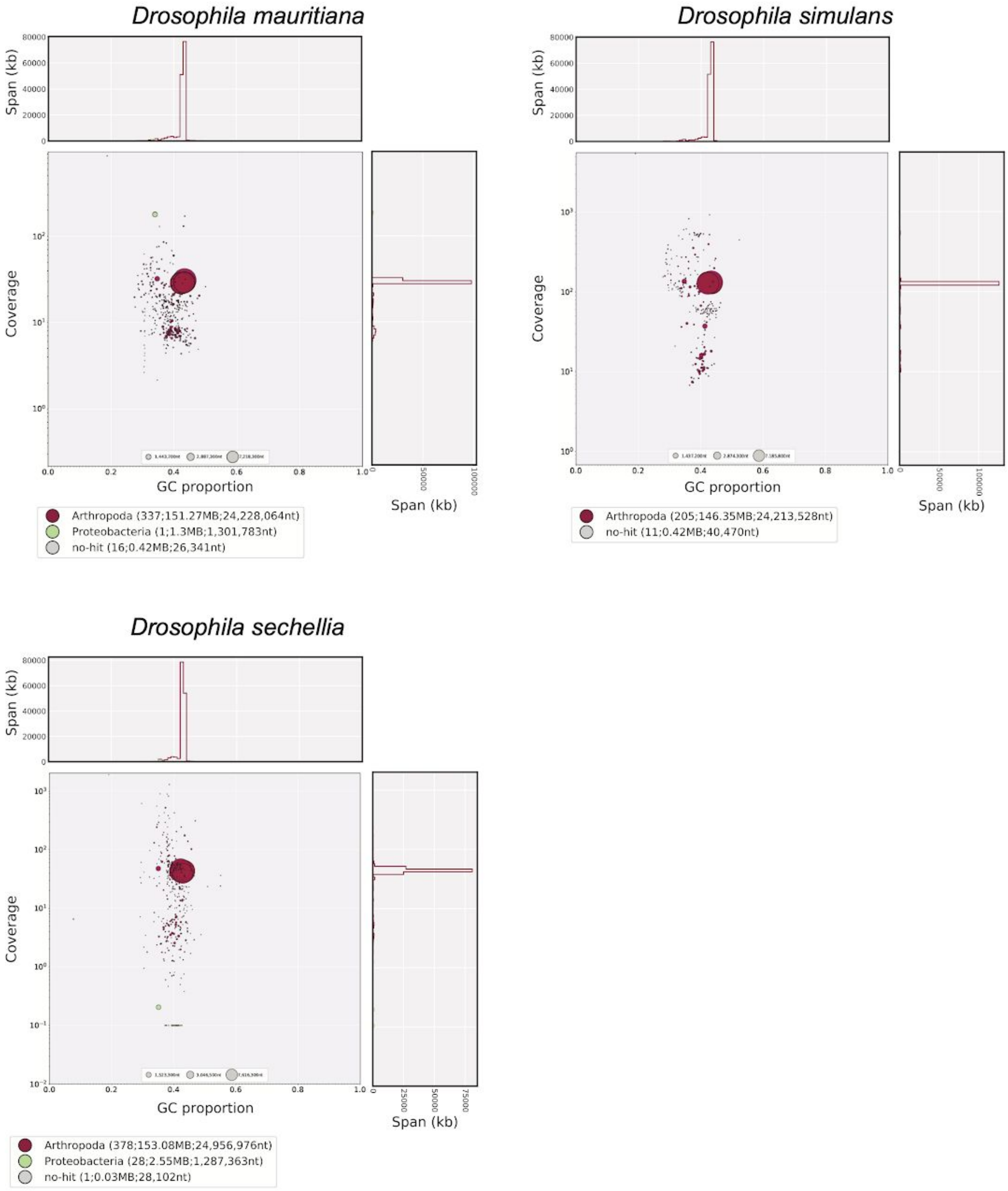


Figure S28. The two-dimensional scatter plot with Illumina coverage and GC histogram (blotplot) in the sim-complex. We used BLAST+ v2.6.0 (Altschul et al. 1990) with blobtools (0.9.19.4; (Laetsch and Blaxter 2017)) to search the nt NCBI database (parameters “-task megablast -max_target_seqs 1

-max_hsps 1 -evalue 1e-25") and assign the homology of each contig. We calculated the Illumina coverage of each contig using SRR483621, SRR8247551, and SRR9030358 for *D. mauritiana*, *D. simulans* and *D. sechellia*, respectively.

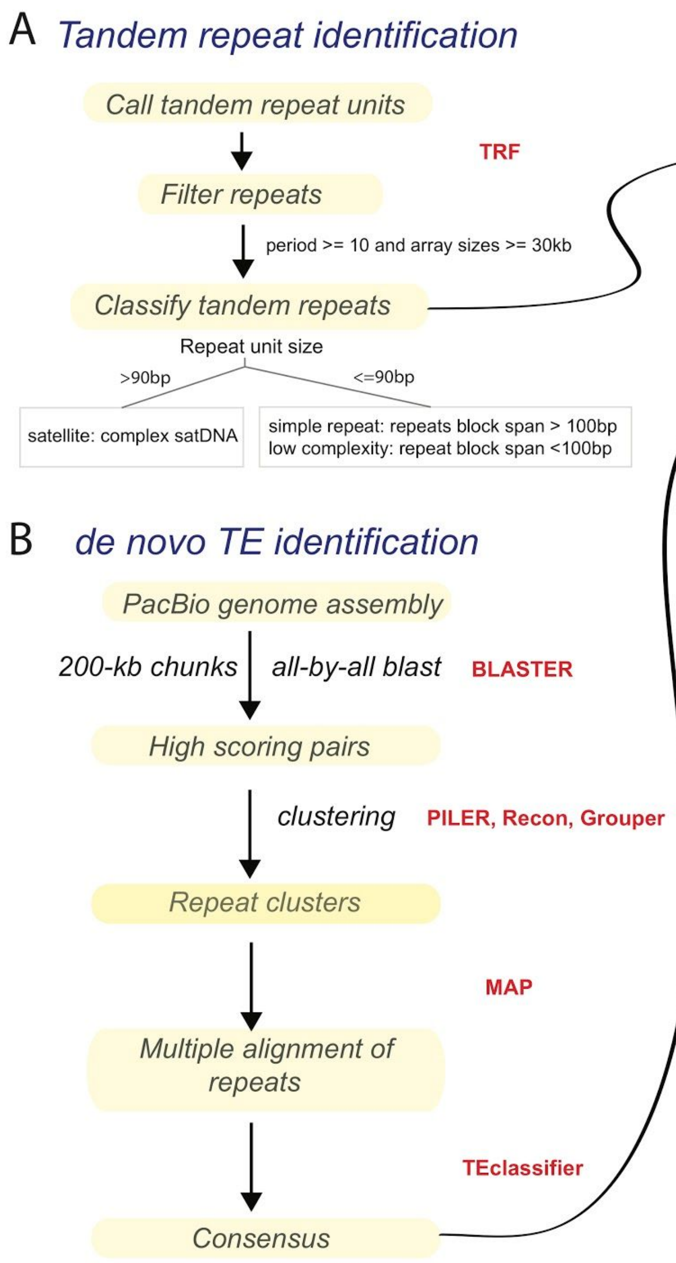


Figure S29.The overview of our repeat annotation approach. We constructed a custom repeat library by combining the Repbase release (20150807) for *Drosophila* with complex satellites with repeat units >90 bp and novel TEs. (A) We identified unannotated complex satellites using Tandem Repeat Finder (TRF) as predicted repeats with period >= 10 and array sizes >= 30kb. (B) We also annotated novel TEs in the sim-complex species using the REPET TE annotation package (Flutre et al. 2011), which includes both *de novo* (Grouper, Piler, and Recon) and homology-based TE annotation programs (RepeatMasker, Censor). We first fragmented the genome sequence into multiple 200kb sequences. These fragments were aligned to themselves using BLAST (Altschul et al. 1990) and clustered using Recon, Grouper and/or Piler (Bao and Eddy 2002; Edgar and Myers 2005). (C) We removed redundant and previously annotated repeats found in Repbase release and the NR database from the Drosophila Repbase release (20150807) and then added newly annotated complex satellite and TEs. We updated some repeat classifications, categorizing some repeats as ‘other’, MINIME as ‘non-LTR’, and the subtelomeric HETRP as ‘satellite’. We used the resulting library (Supplemental File S1) to annotate the three sim-complex species and the *D. melanogaster* reference with RepeatMasker v4.0.5.

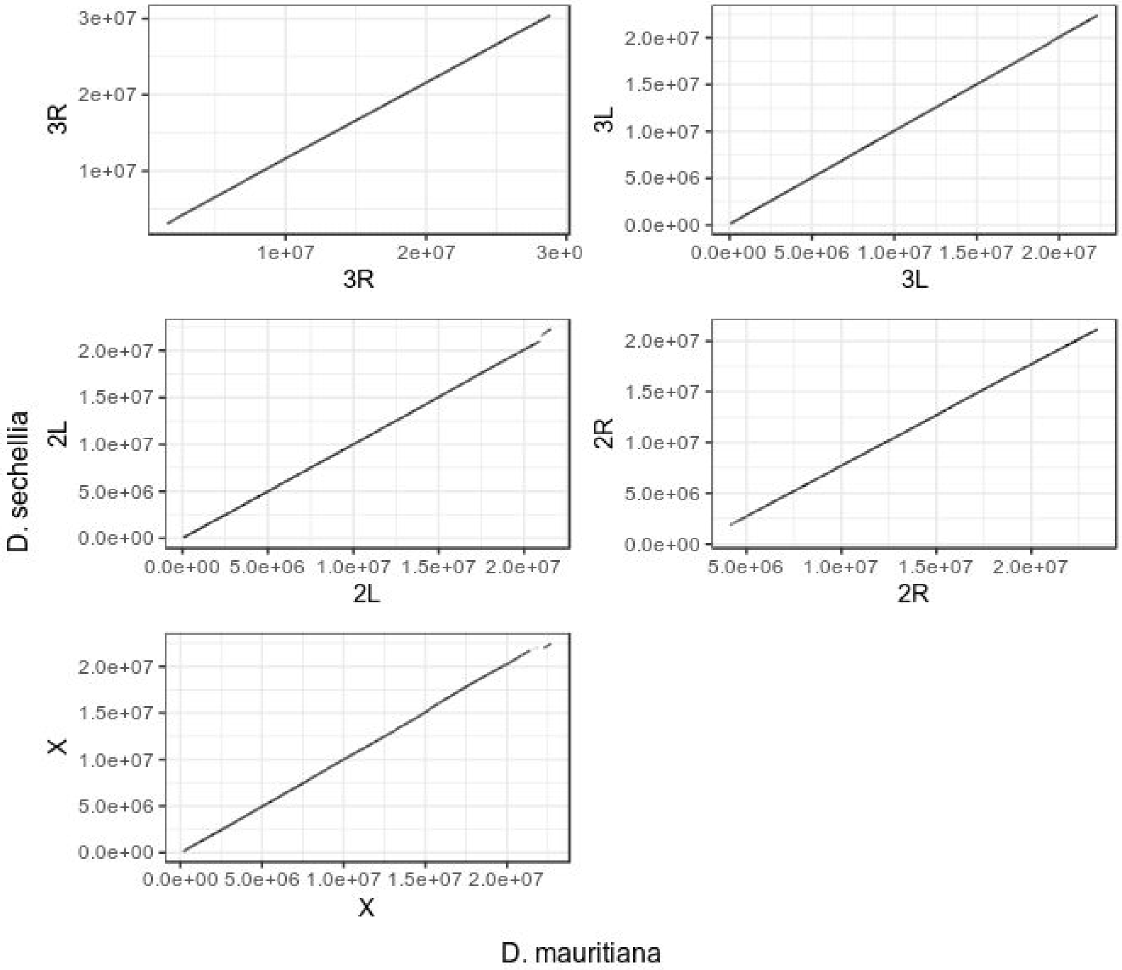


Figure S30. Dotplot of the syntenic alignments that were used in shared TE analysis between *D. mauritiana* and *D. sechellia*. Any annotated TE present within these syntenic alignments were considered to be shared between the two species. The same pipeline was used for the other species pairs (mau-sim, mau-mel).

### Supplementary tables

Table S1. The coverage of PacBio reads for the sim-complex species genomes. We calculated the coverage in 10-kb windows on autosomes, X, and Y chromosomes, respectively. We partitioned the coverage of euchromatic regions (A and X) and heterochromatic regions (Ahet and Xhet) to identify the potential sequencing bias.

Table S2. Contiguity of assemblies reported here compared to that of the *Drosophila melanogaster* community assembly. The assembly size is the total number of basepairs reported across all sequences for the specified assembly. Contig assemblies represent contiguous gapless sequences whereas scaffold assemblies represent collections of contigs ordered and oriented appropriately, but with gaps connecting them (see Methods). NGX is a measure of contiguity indicating that sequences of this length or longer comprise X% of a specified genome size, G. LGX is a measure of contiguity indicating the smallest number of sequences required to represent X% or more of the genome. For this study, the specified genome size, G, is 150 Mb.

Table S3. Quast results of assemblies from the Illumina mapping. We evaluate the quality of assemblies based on the mapping status of Illumina data. Our assemblies improve the total mapping rate by 0.5–2% and properly-paired ratio rate by 5–10% compared to the previously published assemblies. Female reads cover less of the genome for the assemblies presented here than previous assemblies because the Y-chromosome is absent in previous work.

Table S4. Estimates of the nucleotide accuracy of the sim-complex assemblies. Following a previously described method (Koren et al. 2017; Solares et al. 2018), we used Illumina reads to estimate the error rate of nucleotides in our assembly. QV represented as $-10\log_{10}E/T$, where E is the sum of total bases changed (added, deleted, substituted) and T is the total number of bases (minimum coverage of 3).

Table S5. The completeness of our and existing assemblies as assessed by BUSCOs (tab BUSCO summary) and the locations of the duplicated BUSCOs in the alternate contigs of the *D. simulans* assembly (tab D. simulans althap BUSCOs). In the BUSCO summary, we report the number of complete, duplicated, fragmental and missing genes in 2799 conserved BUSCO genes from Diptera.

Table S6. The number of orthologous genes in the annotation of previous assemblies and our assemblies.

Table S7. The redundant contigs from heterozygosity and contigs from the *Drosophila* symbionts. We used Masurca to detect redundant contigs as smaller contigs greater than 40 kb with >90% identity, or between 10 and 40 kb with >95% identity, to the longer contigs. The contigs from the *Drosophila* symbionts are selected by BLAST with blobtools (Laetsch and Blaxter 2017) to search the NCBI nt database.

Table S8. The number of genomic rearrangements in euchromatic and heterochromatic regions of each chromosome arm. We aligned the genomes of the sim-complex and *D. melanogaster* using the mauve. The number of genome rearrangements is calculated by the DCJ model implemented in mauve. The numbers listed above diagonal are rearrangements in heterochromatic regions and the numbers below diagonal are those in euchromatic regions. The Chromosome 4 is presented as a whole because it has an unconventional heterochromatic structure.

Table S9. Percentage of total euchromatic bases comprised of satellite DNA for the autosomes (averaged across autosomal arms) and the X chromosome, and the fold enrichment of satellites on the X chromosome relative to the autosomes. Enrichment was calculated by taking % on the X chromosome divided by % on autosomes.

Table S10. The intron size in *D. melanogaster* and the sim-complex species. Tabulations are homologous intron lengths based on the intron positions (i.e. whole genome, euchromatin, heterochromatin, or the dot chromosome) and intron TE composition (i.e. simple or complex) for each species.

Table S11. List of duplicates that are present in all three sim-complex species but absent in *D. melanogaster* reference strain (ISO1 R6). The first three columns represent the coordinates of the *D. melanogaster* sequence that is duplicated in the sim-complex species. The final three columns correspond to the coordinates in *D. simulans*.

Table S12. Counts of all tRNAs annotated by isotype and predicted anticodon. Counts of tRNAs predicted to be pseudogenes are shown in parentheses.

Table S13. The copy number of each exon in 11 conserved Y-linked genes in assemblies. We used BLAST to detect the homology of each exon. The numbers from *D. melanogaster* are from (Chang and Larracuente 2019; Danilevskaya et al. 1991; Usakin et al. 2005). The numbers from the other species are from the sim-complex assemblies described in this manuscript.

Table S14. The information of PCR primers used to validate the duplication of conserved Y-linked genes and the recently Y-linked duplication from other chromosomes, and the 193XP probe used to detect satellites in cytology in the sim-complex.

Table S15. Y-linked duplications derived from other chromosomes in the sim-complex. We detected duplications and calculated their copy numbers by blast and validated the duplication by PCR. We inferred the duplication mechanism by the presence of introns.

Table S16. The manually corrected misassemblies in the sim-complex species.

Table S17. The accession number of sequencing datasets, including DNA-seq and RNA-seq, and assemblies generated and used in this study.

Table S18. The mapping summary of RNA-seq reads for annotation and transcriptomic analyses in the sim-complex.
